# Population Structure, Stratification and Introgression of Human Structural Variation

**DOI:** 10.1101/746172

**Authors:** Mohamed A. Almarri, Anders Bergström, Javier Prado-Martinez, Fengtang Yang, Beiyuan Fu, Alistair S. Dunham, Yuan Chen, Matthew E. Hurles, Chris Tyler-Smith, Yali Xue

## Abstract

Structural variants contribute substantially to genetic diversity and are important evolutionarily and medically, yet are still understudied. Here, we present a comprehensive analysis of deletions, duplications, insertions, inversions and non-reference unique insertions in the Human Genome Diversity Project (HGDP-CEPH) panel, a high-coverage dataset of 911 samples from 54 diverse worldwide populations. We identify in total 126,018 structural variants (25,588 <100 bp in size), of which 78% are novel. Some reach high frequency and are private to continental groups or even individual populations, including a deletion in the maltase-glucoamylase gene *MGAM* involved in starch digestion, in the South American Karitiana and a deletion in the Central African Mbuti in *SIGLEC5,* potentially leading to immune hyperactivity. We discover a dynamic range of copy number expansions and find cases of regionally-restricted runaway duplications, for example, 18 copies near the olfactory receptor *OR7D2* in East Asia and in the clinically-relevant *HCAR2* in Central Asia. We identify highly-stratified putatively introgressed variants from Neanderthals or Denisovans, some of which, like a deletion within *AQR* in Papuans, are almost fixed in individual populations. Finally, by *de novo* assembly of 25 genomes using linked-read sequencing we discover 1631 breakpoint-resolved unique insertions, in aggregate accounting for 1.9 Mb of sequence absent from the GRCh38 reference. These insertions show population structure and some reside in functional regions, illustrating the limitation of a single human reference and the need for high-quality genomes from diverse populations to fully discover and understand human genetic variation.

## Introduction

Despite the progress in sampling many populations, human genomics research is still not fully reflective of the diversity found globally (Sirugo et al., 2019). Understudied populations limit our knowledge of genetic variation and population history, and their inclusion is needed to ensure they benefit from future developments in genomic medicine. Whole-genome sequencing projects have provided unprecedented insights into the evolutionary history of our species; however, they have mostly concentrated on substitutions at individual sites, although structural variants (affecting > 50bp), which include deletions, duplications, inversions and insertions, contribute a greater diversity at the nucleotide level than any other class of variation and are important in genome evolution and disease susceptibility (Huddleston & Eichler 2016).

Previous studies surveying global population structural variation have examined metropolitan populations at low-coverage (Sudmant et al., 2015a), or a few samples from a larger number of populations (Sudmant et al., 2015b), allowing broad continental comparisons but limiting detailed analysis within each continental group and population. In this study, we present the structural variation analysis of the Human Genome Diversity Project (HGDP)-CEPH panel (Figure 1A), a dataset composed of 911 samples from 54 populations of linguistic, anthropological and evolutionary interest (Cann et al., 2002). We generate a comprehensive resource of structural variants from these diverse and understudied populations, explore the structure of different classes of structural variation, characterize regional and population-specific variants and expansions, discover putatively introgressed variants and identify sequences missing from the GRCh38 reference.

**Figure 1:**
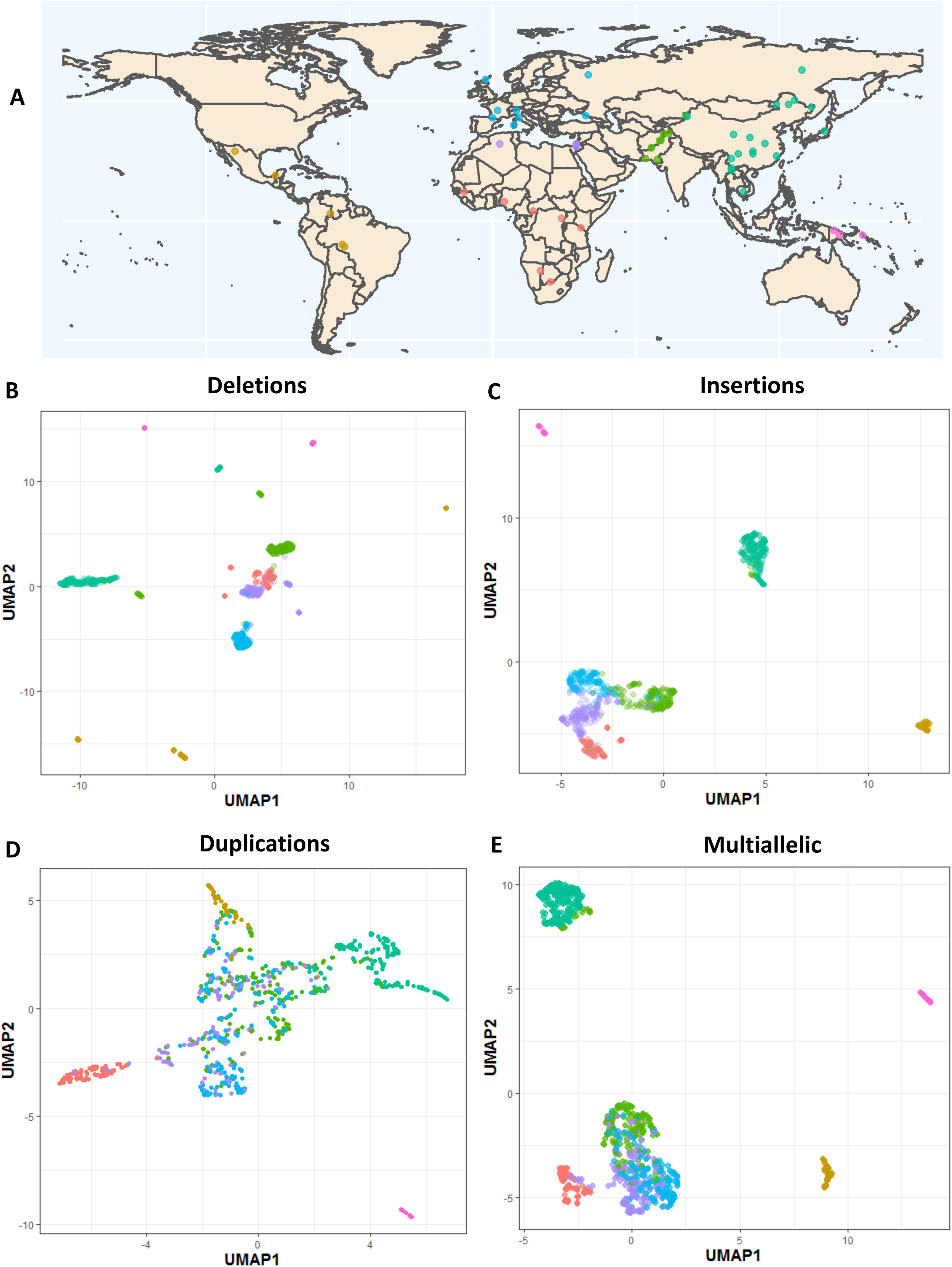
The HGDP dataset and population structure. **A:** The HGDP dataset, each point and colour represents a population and its regional label, respectively. Colours of regional groups are consistent throughout the study. See Table S1 for more details. **B:** UMAP of biallelic deletions genotypes. See Figure S6 for more details. **C:** UMAP of insertions. **D:** UMAP of biallelic duplications. **E:** UMAP of multiallelic variants.

## Results

### Variant Discovery and Comparison with Published Datasets

We generated 911 whole-genome sequences at an average depth of 36x and mapped reads to the GRCh38 reference (Bergström et al., 2019). As the dataset was generated from lymphoblastoid cell lines, we searched for potential cell-line artefacts by analysing coverage across the genome and excluded samples containing multiple aneuploidies, while masking regions which show more limited aberrations (Figure S1). We find many more gains of chromosomes than losses, and in agreement with a previous cell-line based study (Redon et al. 2006), we observe that most trisomies seem to affect chromosomes 9 and 12, suggesting that they contain sequences that enhance proliferation once duplicated in culture. Nevertheless, these cell line artefacts can readily be recognised, and are excluded from the results below.

We identified 126,018 structural variants relative to the reference. These included 25,588 (∼20% of the total) that are smaller than 100bp. We compared our dataset to published structural variation catalogues (Sudmant, et al. 2015a; Sudmant, et al. 2015b), and find that ∼78% of the variants identified in our dataset are not present in the previous studies. Despite having a smaller sample size compared to the 1000 Genomes phase 3 release (Sudmant, et al. 2015a), we discover a higher total number of variants across all different classes of variants investigated. These novel calls are not limited to rare variants, as a considerable number of common and even high-frequency variants are found in regional groups and individual populations (Figure S9). The increased sensitivity reflects the higher coverage, longer reads, improved discovery tools and the large number of diverse populations in our study. Notably, our resource identifies the abundant, but understudied class of small variants (50bp – 100bp), which were not particularly characterized by the Simons Genome Diversity Project (Sudmant, et al. 2015b). At this size range, ∼91% of variants in our dataset are not present in either published catalogues. Collectively, this illustrates that a substantial amount of global structural variation was previously undocumented, emphasizing the importance of studying underrepresented human populations.

### Population Structure

A uniform manifold approximation and projection (UMAP) of deletion genotypes shows clear separation of continental groups, and in many cases even individual populations are distinguished (Figure 1B). Deeply divergent African populations such as the Mbuti, Biaka and San form their own clusters away from the rest of the African populations; admixed groups such as the Hazara and Uygur cluster separately from the Central & South Asian and East Asian groups, while drifted populations such as the Kalash in addition to American and Oceanian populations are clearly differentiated. For less clearly defined populations projecting into continental clusters, we observe examples of finer structure with samples from individual populations appearing closer to themselves relative to other groups (Figure S6).

Insertions, duplications, multiallelic variants and inversions also show some degree of population structure, although less defined in comparison to deletions (Figures 1C-E and S4). Strikingly, the Oceanian populations always remain well-differentiated. Consequently, we find that all classes of genetic variation show population structure, with the observed differences likely reflecting the varying mutational patterns generating each class of structural variant, in addition to the overall number of discovered variants in each class.

### Population Stratification and Selection

Selective pressures can result in highly stratified variants between populations. We assessed the relationship between average population differentiation and the maximal variant allele frequency difference for each population pair (Figure 2A-C). Outliers in this relationship, i.e. variants that show a higher allele frequency difference than expected, have been proposed to be under selection (Coop et al., 2009; Huerta-Sanchez et al., 2014). Both deletions and insertions show similar distributions, while biallelic duplications display lower stratification. We do see some notable outliers, for example the Lowland/Sepik Papuans are almost fixed (86%) for a deletion in *HBA2*, which is absent in Papuan Highlanders. High frequencies of α-globin deletions have been suggested to be protective against malaria, which is not found in the highlands of Papua New Guinea, but is present in the lowlands (Yenchitsomanus et al., 1985, Flint et al., 1986). On the other hand, Papuan Highlanders have a small insertion (123bp) near an exon of *VGLL4* at 93% frequency which is markedly less common in Papuan Lowlands (7%). We also find a deletion within *MYO5B* that is particularly common (88%) in the Lahu from China, a population shown to have high numbers of private single nucleotide variants in addition to carrying rare Y-chromosome lineages (Bergström et al., 2019).

**Figure 2:**
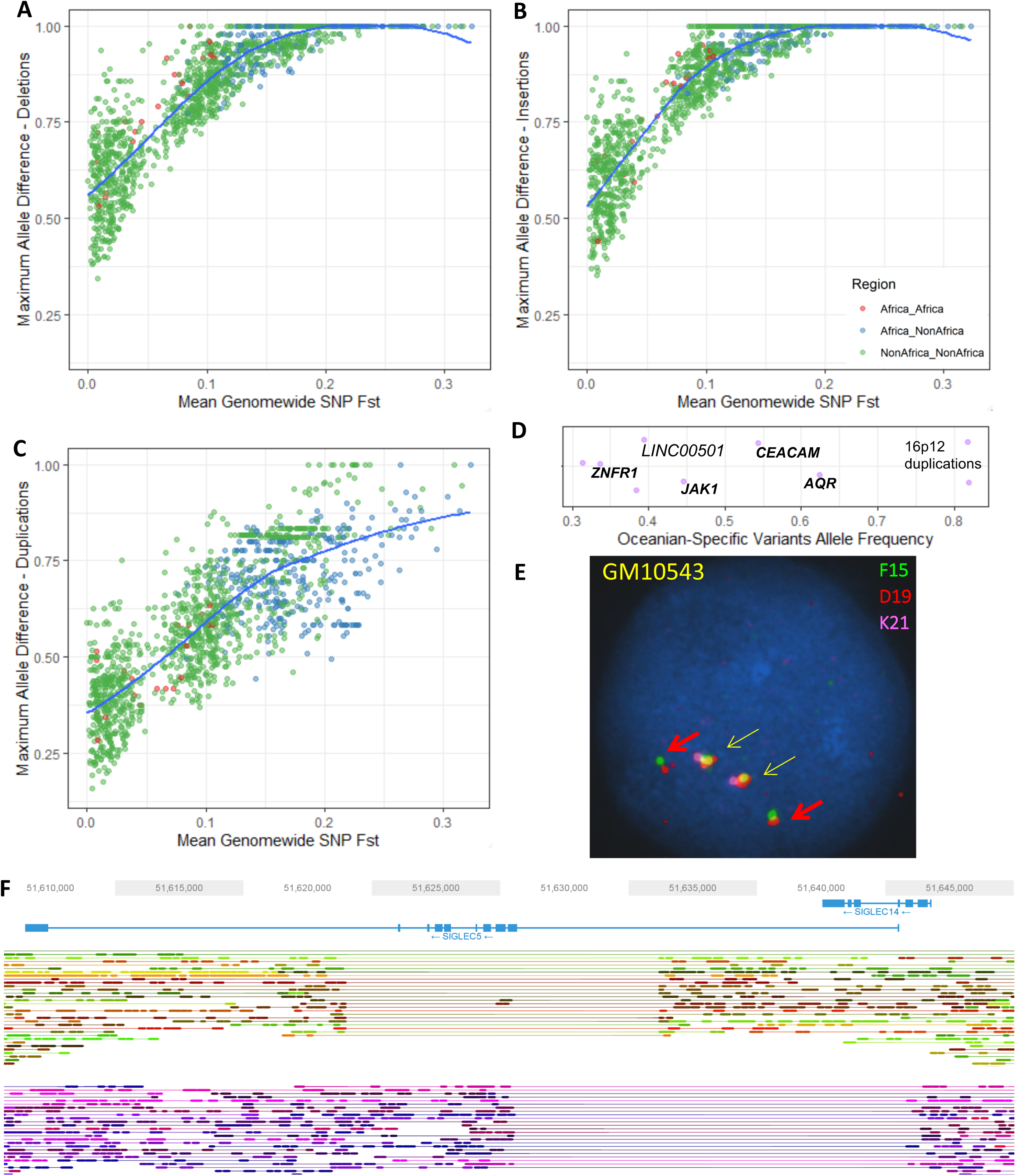
Population stratification of structural variants. **A:** Maximum allele frequency difference of deletions as a function of population differentiation for 1431 pairwise population comparisons. Blue curve represents loess fits. **B-C**: Same as A but for insertion and biallelic duplications, respectively. **D:** High frequency Oceanian-specific variants (>30% frequency). Each point represents a variant with the x-axis illustrating its frequency. Random noise is added to aid visualization. Almost all variants are shared with the Denisovan genome and are within (**bold**) or near the illustrated genes. **E**: Fluorescent in situ hybridization illustrating the 16p12 Oceanian-specific duplication shared with Denisova in a homozygous state (cell-line GM10543). Yellow arrows show reference and red arrow illustrate duplication. See Figure S12-13 for more details. **F**: Distinct deletions at the SIGLEC5/SIGLEC14 locus in an Mbuti sample (HGDP00450) resolved using linked-reads. One haplotype (top) carries the Mbuti-specific variant that deletes most exons in *SIGLEC5* and is present at high frequency (54%), while the second haplotype (bottom) carries a globally common deletion that deletes SIGLEC14, creating a fused gene (See supplementary information for more details).

The large number of samples per population allowed us to investigate population-private variants (Figure S7). We searched for functional effects of such variants and found a 14kb deletion in the South American Karitiana population at 40% frequency. This variant removes the 5’ upstream region of *MGAM* up to the first exon, potentially inactivating the gene which encodes Maltase-glucoamylase, an enzyme highly expressed in the small intestine and involved in the digestion of dietary starches (Nichols et al., 2003). Interestingly, a recent ancient DNA study of South Americans has suggested that selection acted on this gene in ancient Andean individuals, possibly as a result of their transition to agriculture (Lindo et al., 2018). This gene has also been proposed to be under selection in dogs, due to adaptation to a starch-rich diet during domestication (Axelsson et al., 2013). However, the high frequency and presence of individuals homozygous for this deletion suggests that purifying selection on the ability to digest starch has been relaxed in the history of the Karitiana.

We discovered a deletion that is private and at 54% frequency in the Central African Mbuti hunter-gatherer population that deletes *SIGLEC5* without removing its adjacent paired receptor *SIGLEC14* (Figure 2F). Siglecs, a family of cell-surface receptors that are expressed on immune cells, detect sialylated surface proteins expressed on host cells. Most SIGLECs act as inhibitors of leukocyte activation, but *SIGLEC14* is an activating member which is thought to have evolved by gene conversion from *SIGLEC5* (Angata et al., 2006). This evolution has been proposed to result in a selective advantage of combating pathogens that mimic host cells by expressing sialic acids, providing an additional activation pathway (Akkaya and Barclay 2013). The deletion we identify in the Mbuti, however, seems to remove the function of the inhibitory receptor, while keeping the activating receptor intact. This finding is surprising, as paired receptors are thought to have evolved to fine-tune immune responses; and the loss of an inhibitory receptor is hypothesized to result in immune hyperactivity and autoimmune disease (Lübbers et al., 2018).

### Archaic Introgression

We genotyped our calls in the high coverage Neanderthal and Denisovan archaic genomes (Meyer et al., 2012; Prufer et al., 2017; Prufer et al., 2014), and find hundreds of variants that are exclusive to Africans and archaic genomes, suggesting that they were part of the ancestral variation that was lost in the out-of-Africa bottleneck. We then searched for common, highly stratified variants that are shared with archaic genomes but are not present in Africa, possibly resulting from adaptive introgression. We identify variants across a wide range of sizes, the smallest 63bp and largest 30kb (Table 1), and note that almost all lie within or near genes, potentially having functional consequences.

**Table 1:** Allele frequencies of regionally stratified variants shared with high coverage archaic genomes but not found in African populations. Neanderthal refers to both published high coverage genomes. If a variant lies within or intersects gene it is highlighted in bold, otherwise the nearest gene is presented. The deletion within *ALLC* is only shared with the Vindija Neanderthal. The *TNFRSF10D* duplication common in Oceania is also present at low frequency (5%) in Africa. Africans do not have both deletion and duplication variants, which are in linkage disequilibrium in Oceanians (r^2^ = 0.48). The duplications at chr16p12.2 at high frequency in Oceania (82%) are part of a complex structural variant (Figure S12-13). EA - East Asia, ME - Middle East, AMR - America, CSA - Central South Asia, OCE - Oceania.

| Position | Size (bp) | Variant | EUR | CSA | EA | ME | AMR | OCE | Gene | Neanderthal | Denisova |
| --- | --- | --- | --- | --- | --- | --- | --- | --- | --- | --- | --- |
| chr1: 64992619-64992994 | 375 | DEL | 0 | 0 | 0 | 0 | 0 | 0.44 | **JAK1** | REF | DEL |
| chr2:3684113-3690212 | 6099 | DEL | 0.02 | 0.003 | 0.05 | 0.03 | 0 | 0.26 | **ALLC** | DEL<sub>Vin</sub> | REF |
| chr3:177287011-177292441 | 5430 | DEL | 0 | 0 | 0 | 0 | 0 | 0.39 | **LINC00501** | REF | DEL |
| chr8:23124835-23130567 | 5732 | DEL | 0 | 0.02 | 0.002 | 0 | 0 | 0.36 | **TNFRSF10D** | DEL | REF |
| chr8:23134649-23164796 | 30147 | DUP | 0 | 0 | 0 | 0 | 0 | 0.48 | **TNFRSF10D** | DUP | DUP |
| chr11:60460681-60461880 | 1199 | DEL | 0 | 0 | 0.02 | 0 | 0.17 | 0 | **MS4A1** | DEL | REF |
| chr12:101882163-101883377 | 1214 | DEL | 0.02 | 0.08 | 0.32 | 0.007 | 0.01 | 0.33 | **DRAM1** | DEL | REF |
| chr12:104799951-104803150 | 3199 | DUP | 0.003 | 0.009 | 0 | 0.01 | 0 | 0.33 | **SLC41A2** | DUP | REF |
| chr15:34920811-34925992 | 5181 | DEL | 0 | 0 | 0 | 0 | 0 | 0.63 | **AQR** | REF | DEL |
| chr16p12.2 | Complex | DUP | 0 | 0 | 0 | 0 | 0 | 0.82 | Multiple | REF | DUP |
| chr16:75059992-75060055 | 63 | DEL | 0 | 0 | 0 | 0 | 0 | 0.34 | **ZNRF1** | DEL | DEL |
| chr17:3038851-3041981 | 3130 | DEL | 0 | 0 | 0 | 0 | 0 | 0.16 | **RAP1GAP2** | DEL | DEL |
| chr19:42529806-42531042 | 1236 | DEL | 0 | 0 | 0 | 0 | 0 | 0.54 | **CEACAM1** | DEL | DEL |

We replicated the putatively Denisovan introgressed duplication at chromosome 16p12.2 exclusive to Oceanians (Sudmant et al. 2015b). We explored the frequency of this variant in our expanded dataset within each Oceanian population, and despite all the Bouganville Islanders having significant East Asian admixture, which is not found in the Papuan Highlanders, we do not find a dilution of this variant in the former population: it is present at a remarkable and similar frequency in all three Oceanian populations (∼82%). These duplications form the most extreme regional-specific variants (Figure 2D, Figure S8), and their unusual allele distribution suggests that they may have remained at high frequencies after archaic introgression due to positive selection. We characterized this variant in more detail using fluorescent in situ hybridization (Figure 2E, Figure S12-13), and find that it consists of a region of the reference sequence that has duplicated and inserted into a gene-rich region ∼7Mb away in chr16p11.2, confirming a recent study (Hsieh et al., 2019). The selective pressure acting on this duplication and its target remain unknown and require further study; however, its similar frequency across the Oceanian populations examined contrasts with the differing frequency of the malaria-associated *HBA2* deletion across Oceania, suggesting that malaria infection is unlikely to be driving the signal we see at the 16p12.2 duplication.

We discover multiple additional high-frequency Oceanian-private variants that are shared with the Denisovan genome (Figure 2D), illustrating the separate introgression event in Oceanians and their subsequent isolation (Browning et al., 2018). A deletion within *AQR*, an RNA helicase gene, is present at 63% frequency and shared only with the Altai Denisovan (Figure S15). The highest expression of this gene is in EBV-transformed lymphocytes (GTEx Consortium, 2013). RNA helicases play an important role in the detection of viral RNAs and mediating the antiviral immune response, in addition to being necessary host factors for viral replication (Ranji & Boris-Lawrie, 2010). AQR has been reported to be involved in the recognition and silencing of transposable elements (Akay et al., 2017), and is known to regulate HIV-1 DNA integration (Konig et al., 2008). Two other notable Denisovan-shared deletions of high frequency are in *JAK1*, encoding a kinase important in cytokine signalling (44%) and *CEACAM1* (also known as CD66a) a glycoprotein part of the immunoglobulin superfamily (54%).

In the Americas we identify a deletion, shared only with Neanderthals, that reaches ∼26% frequency in both the Surui and Pima. This variant removes an exon in *MS4A1* (Figure S16), a gene encoding the B-cell differentiation antigen CD20, which plays a key role in T cell–independent antibody responses and is the target of multiple recently developed monoclonal antibodies for B-cell associated leukemias, lymphomas and autoimmune diseases (Kuijpers et al., 2010; Marshall et al., 2017). This deletion raises the possibility that therapies developed in one ethnic background might not be effective in others, and that access to individual genome sequences could guide therapy choice.

Both Neanderthals and Denisovans thus appear to have contributed potentially functional structural variants to different modern human populations. As many of the identified variants are involved in immune processes (Table 1), it is tempting to speculate that they are associated with adaptation to pathogens after modern humans expanded into new environments outside of Africa.

### Multiallelic Variants and Runaway Duplications

We found a dynamic range of expansion in copy numbers, with variants previously found to be biallelic containing additional copies in our more diverse dataset. Among these multiallelic copy number variants, we find intriguing examples of ‘runaway duplications’ (Handsaker et al., 2015), variants that are mostly at low copy numbers globally, but have expanded to high copy numbers in certain populations, possibly in response to regionally-restricted selection events (Figure 3).

**Figure 3:**
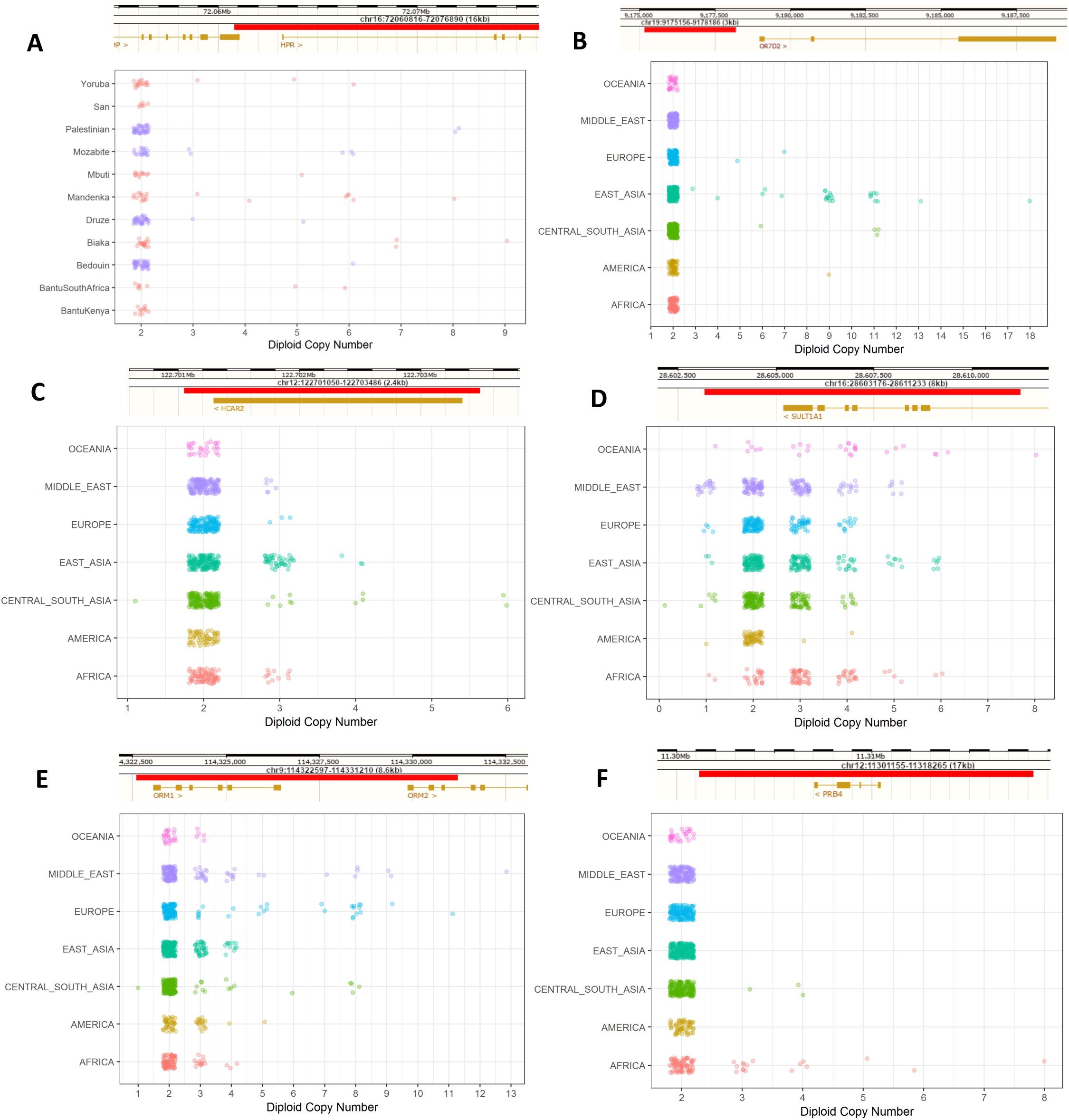
Copy Number Expansions and Runaway Duplications. Red bar illustrates the location of the expansion. Additional examples are shown in Figure S11. **A:** Expansion in *HPR* in Africans and Middle Eastern samples. **B:** Expansions upstream *OR7D2* that are mostly restricted to East Asia. The observed expansions in Central & South Asian samples are all in Hazara samples, an admixed population carrying East Asian ancestry. **C:** Expansions within *HCAR2* which are particularly common in the Kalash population. **D:** Expansions in *SULT1A1* which are pronounced in Oceanians (median copy number, 4; all other non-African continental groups, 2; Africa, 3). **E:** Expansions in *ORM1/ORM2*. This expansion was reported previously in Europeans (Handsaker et al., 2015); however, we find it in all regional groups and particularly in Middle Eastern populations. **F:** Expansions in *PRB4* which are restricted to Africa and Central & South Asian samples with significant African admixture (Makrani and Sindhi).

We discover multiple expansions that are mostly restricted to African populations. The hunter-gatherer Biaka are notable for a private expansion downstream of *TNFRSF1B* that reaches up to 9 copies (Figure S11). We replicated the previously identified *HPR* expansions (Figure 3A), and find that they are present in almost all African populations in our study (Handsaker et al., 2015, Sudmant et al., 2015b). *HPR* encodes a haptoglobin-related protein associated with defense against trypanosome infections (Smith et al., 1995). We observe populations with the highest copy numbers to be Central and West African, consistent with the geographic distribution of the infection (Franco et al., 2014). In contrast to previous studies, we also find the expansion at lower frequencies in all Middle Eastern populations, which we hypothesize is due to recent gene flow from African populations.

We identified a remarkable expansion upstream of the olfactory receptor *OR7D2* that is almost restricted to East Asia (Figure 3B), where it reaches up to 18 copies. Haplotype phasing demonstrates that many individuals contain the expansion on just one chromosome, illustrating that these alleles have mutated repeatedly on the same haplotype background. However, we identify a Han Chinese sample that has a particularly high copy number. This individual has nine copies on each chromosome, suggesting that the same expanded runaway haplotype is present twice in a single individual. This could potentially lead to an even further increase in copy number due to non-allelic homologous recombination (Handsaker et al., 2015).

We discovered expansions in *HCAR2* (encoding HCA_2_) in Asians which are especially prominent in the Kalash group (Figure 3C), with almost a third of the population displaying an increase in copy number. HCA_2_ is a receptor highly expressed on adipocytes and immune cells, and has been proposed as a potential therapeutic target due to its key role in mediating anti-inflammatory effects in multiple tissues and diseases (Offermanns 2017). Another clinically-relevant expansion is in *SULT1A1* (Figure 3D), which encodes a sulfotransferase involved in the metabolism of drugs and hormones (Hebbring et al., 2008). Although the copy number is polymorphic in all continental groups, the expansion is more pronounced in Oceanians.

### *De novo* assemblies and sequences missing from the reference

We sequenced 25 samples from 13 populations using linked-read sequencing at an average depth of ∼50x and generated *de novo* assemblies using the Supernova assembler (Weisenfeld et al., 2017) (Table S2). By comparing our assemblies to the GRCh38 reference, we identified 1631 breakpoint-resolved unique, non-repetitive insertions across all chromosomes which in aggregate account for 1.9Mb of sequences missing from the reference (Figure 4A). A San individual contained the largest number of insertions, consistent with their high divergence from other populations. However, we note that the number of identified insertions is correlated with the assembly size and quality (Figure S18), suggesting there are still additional insertions to be discovered.

**Figure 4:**
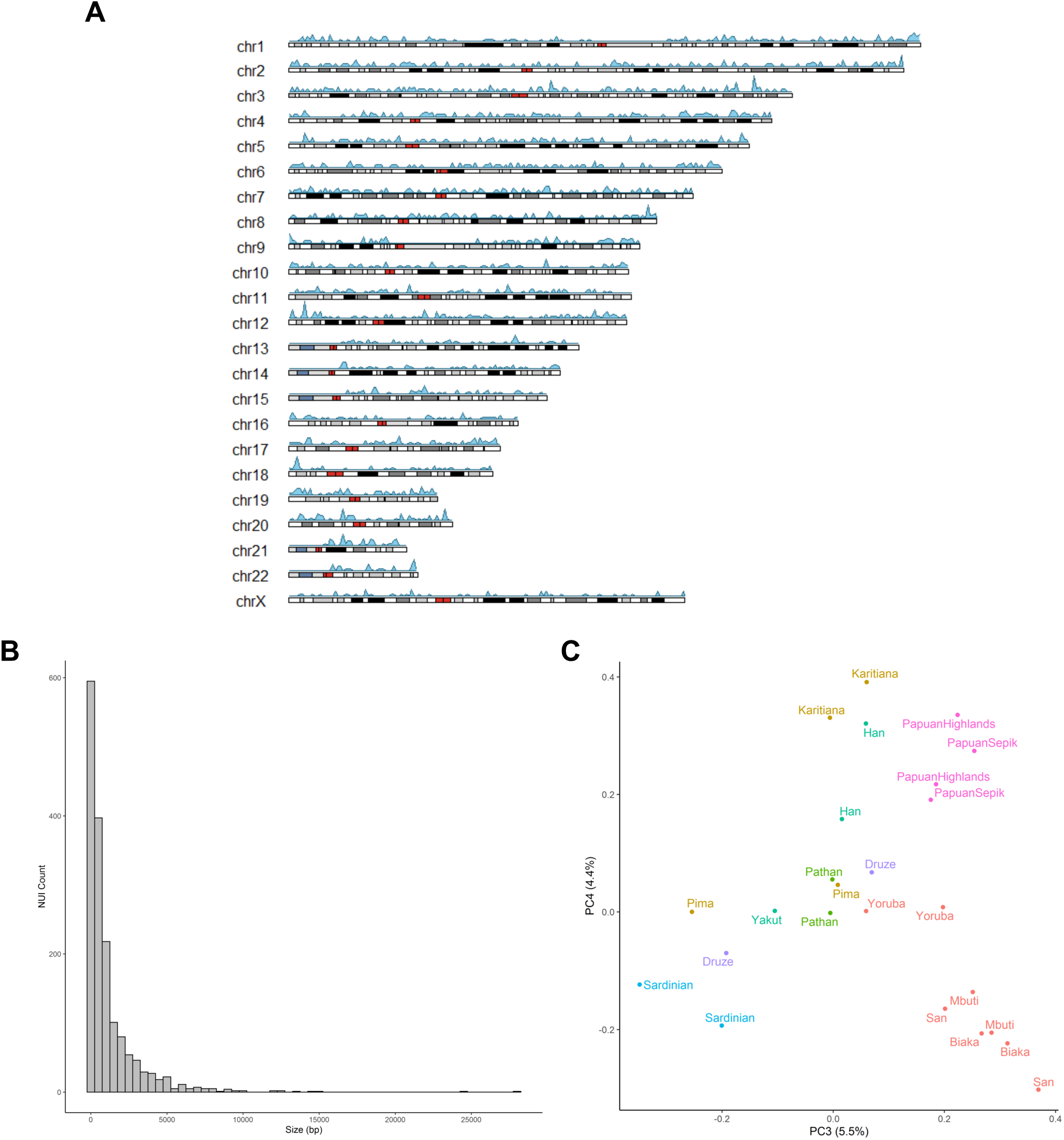
Non-Reference Unique Insertions (NUIs). **A:** Ideogram illustrating the density of identified NUI locations across different chromosomes using a window size of 1 Mb. Colours on chromosomes reflect chromosomal bands with red for centromeres. **B**: Size distribution of NUIs using a bin size of 500bp. **C**: PCA of NUI genotypes showing population structure (PC3-4). Previous PCs potentially reflect variation in size and quality of the assemblies.

We find that the majority of insertions are relatively small, with a median length of 513bp (Figure 4B). They are of potential functional consequence as 10 appear to reside in exons. These genes are involved in diverse cellular processes, including immunity (*NCF4*), regulation of glucose (*FGF21*), and a potential tumour suppressor (*MCC*). Although many insertions are rare - 41% are found in only one or two individuals - we observe that 290 are present in over half of the samples, suggesting the reference genome may harbour rare deletion alleles at these sites. These variants show population structure, with Central Africans and Oceanians showing most differentiation (Figure 4C), reflecting the deep divergences within Africa and the effect of drift, isolation and possibly Denisovan introgression in Oceania. While the number of *de novo* assembled genomes using linked or long reads is increasing, they are mostly representative of urban populations. Here, we present a resource containing a diverse set of assemblies with no access or analysis restrictions.

## Discussion

In this study we present a comprehensive catalogue of structural variants from a diverse set of human populations. Our analysis illustrates that a substantial amount of variation, some of which reaches high frequency in certain populations, has not been documented in previous sequencing projects. The relatively large number of high-coverage genomes in each population allowed us to identify and estimate the frequency of population-specific variants, providing insights into potentially geographically-localized selection events, although further functional work is needed to elucidate their effect. Our finding of common clinically-relevant regionally private variants, some of which appears to be introgressed from archaic hominins, argues for further efforts generating genome sequences without data restrictions from under-represented populations. We note that despite the diversity found in the HGDP panel, considerable geographic gaps remain in Africa, the Americas and Australasia.

The use of short reads in this study restricts the discovery of complex structural variants, demonstrated by recent reports which uncovered a substantially higher number of variants per individual using long-read or multi-platform technologies (Audano et al., 2019; Chaisson et al., 2019). Additionally, comparison with a mostly linear human reference formed from a composite of a few individuals, and mainly from just one person, limits accurately representing the diversity and analysis of human structural variation (Schneider et al., 2017). The identification of considerable amounts of sequences missing from the reference, in this study and others (Wong et al., 2018; Sherman et al., 2019), argues for the creation of a graph-based pan-genome that can integrate structural variation (Garrison et al., 2018). Such computational methods and further developments in long-range technologies will allow the full spectrum of human structural variation to be investigated.

## Data availability

Raw read alignments are available from the European Nucleotide Archive (ENA) under study accession number PRJEB6463. The 10x Genomics linked-reads data are available at ENA under study accession PRJEB14173. Structural variant calls, Supernova *de novo* assemblies and NUI fastas are available on ftp://ngs.sanger.ac.uk/scratch/project/team19/HGDP_SV/

## Acknowledgments

We thank Richard Durbin, Hélène Blanché, Thomaz Pinotti, Klaudia Walter and members of the Tyler-Smith group for advice and discussions. We also thank Robert Handsaker for technical advice on the structural variant discovery algorithm. We would particularly like to thank all the individuals who donated or collected samples for this study and the CEPH Biobank at Fondation Jean Dausset-CEPH for the maintenance and distribution of the HGDP DNAs. M.A.A., A.B., J.P.-M., A.S.D., Y.C., C.T.-S and Y.X. were supported by Wellcome grant 098051.

## Author Contributions

Y.X. and C.T.-S. conceived and supervised the study. M.A.A. designed the study and led the analysis with contributions from A.B., J.P.-M., A.S.D and Y.C.

F.Y. designed, performed and interpreted FISH results. B.F. performed FISH, image capture and analysis. M.E.H. assisted interpretation of results. M.A.A. wrote the manuscript with contribution from all authors.

## Materials, Methods and Supplementary Information

### Sample Sequencing and Read Processing

Number of samples per population and regional labels are presented in Table S1. For more detailed information on the population labels, sequencing and mapping process of the samples analysed in this dataset, refer to Bergström et al., 2019. In brief, samples investigated in this project were provided by the HGDP-CEPH (Cann et al., 2002). Ten samples (PCR) were sequenced in a previous study for comparison with the Denisovan genome (Meyer et al., 2012), all using PCR-based libraries (subsequently called “Meyer” samples). An additional 142 samples were sequenced as part of the Simon Genome Diversity Project (“SGDP”), mostly using PCR-free methods (Mallick et al., 2016). The remaining 808 samples were sequenced at the Wellcome Sanger Institute using either library preparation method, and in some cases both on the same sample, resulting in 823 genome sequences (“Sanger” samples). Twelve SGDP and two Meyer samples were also independently sequenced at Sanger. Each of the Sanger, SGDP and Meyer samples used sequencing technologies with different read lengths (2×151 bp, 2×100 bp, and 94+100 bp or 95+101), mean depth (35x, 42.4x, 28x) and insert sizes (447 bp, 310 bp, 264 bp) respectively. All sample reads were processed through the automated pipeline of the Wellcome Sanger Institute sequencing facility and mapped to the GRCh38 reference. Number of samples per population and per library preparation is presented in Table S1.

### Sample Quality Control

As the whole dataset is derived from lymphoblastoid cell lines, we searched for potential cell-line artefacts by analysing local coverage of each sample. Coverage was calculated at ∼300,000 single positions across the genome and a rolling mean was plotted, normalized by the genome-wide median. Each chromosome in all samples was manually inspected for variation in depth.

In the SNP analysis (Bergström et al., 2019), a total of 929 samples remained after quality control, including some samples exhibiting copy number gains, as these were observed to have minor effects on genotyping accuracy. Here, we subsequently excluded an additional 10 samples that show putative cell-line artefacts across multiple chromosomes (Figure S1). For samples showing more limited putative artefacts, we masked such regions and set any calls within them to missing. A total of 74 samples contained masked regions. This included the sex chromosomes, where we identify many instances of partial loss of Y chromosomes in addition to observing a single XXY male, which could be a natural occurrence rather than an artefact. This resulted in a total dataset of 919 samples composed of 644 Sanger PCR-free, 147 Sanger PCR, 111 SGDP PCR-free, 9 SGDP PCR and 8 Meyer.

### Variant Calling and Quality Control

Two recent studies have comprehensively evaluated different short-read structural variant callers and provided recommendations and best practices (Cameron et al., 2019; Kosugi et al., 2019). We choose to use Manta (Chen et al., 2016), an assembly-based caller, as it performed well in these studies. Additionally, we used GenomeSTRiP (Handsaker et al., 2015), which uses read-depth and read-pair information to identify copy number variants as we were interested in multiallelic variants. GenomeSTRiP v2.00 and Manta v1.6 were run using default parameters. GenomeSTRiP identifies deletions, duplications and multiallelic variants > 1kb, while Manta identifies deletions, insertions, inversions, tandem duplications and interchromosomal translocations >50bp.

#### GenomeSTRiP

We first ran the algorithm jointly on all libraries, including libraries not passing quality control for short variant calling. We subsequently found the Meyer libraries to have lower quality of calls and re-ran the algorithm excluding them.

Duplicate samples prepared using both PCR and PCR-free libraries were created for quality control purposes. We ran GenomeSTRiP twice, once including the PCR prepared duplicates, and the second with PCR-free duplicates, together with the rest of the dataset. Comparing both callsets revealed that PCR-based libraries contained a higher number of shared heterozygous calls that were missing from the PCR-free libraries. These calls were excluded using the excessive heterozygosity tag calculated by bcftools v1.9 (ExcHet < 0.0001) separately for each library preparation and sequencing location set (i.e. SGDP PCR, SGDP PCR-free, Sanger PCR and Sanger PCR-free). For the SGDP PCR samples we used ExcHet < 0.05 due to this callset only having 9 samples. After this QC, a VCF with 50,474 CNVs from 911 samples was generated. We find no detectable batch effects, with the top PCs displaying continental variation and subsequently population variation (Figure S4A-B).

We examined the callset and identified instances where the algorithm splits single variants into multiple shorter entries which are not always overlapping. This a known behaviour of the GenomeSTRiP CNV pipeline which seems to occur if a low quality variant is found within a larger CNV or when there are variants with different copy numbers across different individuals within a sub-segment of a larger variant. To more accurately estimate the total number of identified CNVs in our dataset accounting for these issues, we merged high quality (CNQ > 12) calls that have the same diploid copy number and are within 50 kb of each other, for each sample separately. At this step we found one sample (HGDP01254) that, although not showing any observable chromosomal abnormalities, contained slightly elevated numbers of variants compared to the rest of the samples. These calls had relatively low genotype quality. We chose to be conservative and subsequently excluded this sample from the GenomSTRiP callset, leaving 910 individuals. All variants were then merged using bedmap v2.4.35 (Neph at el., 2012) based on 100% overlap. This identified 39,634 autosomal variants, 1,102 variants on the X-chromosome and 289 variants on the Y-chromosome. 22,914 variants were composed of biallelic deletions, 16,012 were duplications, and 2,099 were variants with both deletion and duplication alleles (Figure S2).

#### Manta + Graphtyper2

We ran Manta v1.6 (Chen et al., 2016) on the 911 libraries discussed above to generate individual VCFs for each sample. We then extracted variants that ‘PASS’ all the quality thresholds of the algorithm. In Manta v1.6, inversions are reported as breakends (BND), we subsequently used a script provided with the Manta download (`convertInversion.py`) to convert them into single inverted sequence junctions, as represented in previous versions. We masked the potential cell-line artefact regions identified in the samples as in the previous step. We subsequently merged all samples using svimmer (https://github.com/DecodeGenetics/svimmer) under default conditions, as performed in Eggertsson et al., 2019. The merged dataset comprised 160,958 variants.

As the Manta call set is not joint-called, differences in read lengths, insert sizes, coverage and library preparation in the HGDP dataset may create batch effects. Additionally, a variant found in one sample may be present but missed in another sample due to the differing variables mentioned above. To address this, we discarded the original genotypes identified by Manta for each sample and jointly regenotyped the merged dataset across all samples concurrently using Graphtyper2 (Eggertsson et al., 2019). This algorithm creates an acyclic mathematical graph structure to represent the reference genome and identified structural variants, to which reads are then re-aligned and genotyped. The algorithm provides three different genotyping models: ‘coverage’, ‘breakpoint’ and also an ‘aggregate’ model that uses information from the two previous models. We extracted the ‘aggregate’ model as suggested for all variants (Eggertsson et al., 2019), except inversions which we used the breakpoint model (no aggregate model was identified for inversions). We excluded variants with size over 10 Mb and set all variants with GQ < 20 to missing. We also set variant genotype calls that have a ‘FAIL1’ tag to missing. For duplications, we also so ‘FAIL2’ and ‘FAIL3’ to missing. We excluded variants with (ExcHet < 0.00001) across the entire dataset and also separately for each library and location samples set (SangerPCR < 0.001, SangerPCRfree < 0.00001, SGDP PCR < 0.05, SGDP PCRfree < 0.00001). Finally, we removed any monomorphic variants. To test for batch effects, we ran a principal component analysis separately for each class of variant identified (DEL, DUP, INS, INV). We find no observed batch effects across all classes, with the top PCs displaying continental variation and subsequently population variation (Figure S4C-E). The final analysed Manta callset included 68,089 deletions, 25,084 insertions, 7,290 duplications, 1,895 inversions and 1,667 translocations.

#### Comparison between GenomeSTRiP and Manta+Graphtyper callsets

To identify non-overlapping variants in both callsets we used bedmap v2.4.35 (Neph at el., 2012) with a threshold of 50% reciprocal overlap. This identified 126,018 unique variants. To evaluate the accuracy of genotype calling, we extracted African-specific variants present in both GenomeSTRiP and Manta+Graphtyper callsets. We observe high correlation of variant allele frequencies between both callsets (r = 0.97, Figure S3), with the slight differences partly due to varying missingness.

### Comparison with Published Datasets

To assess the novelty of our dataset, we compared it with two structural variation callsets:

1. The 1000 Genomes Phase 3 Structural Variation Dataset (1000G, Sudmant, et al. 2015a), downloaded from: ftp://ftp.1000genomes.ebi.ac.uk/vol1/ftp/phase3/integrated_sv_map/supporting/GRC h38_positions/
2. The copy number analysis of the Simons Genome Diversity Project (Sudmant, et al. 2015b).

As the SGDP callset is mapped using GRCh37, we used the UCSC LiftOver function to GRCh38 using default parameters (https://genome.ucsc.edu/cgi-bin/hgLiftOver). We observe that 294 variants failed LiftOver and were not further considered. The README file for the downloaded 1000G dataset notes that these variants were lifted over to GRCh38, leading to the exclusion of 121 variants. We lifted over variants from our dataset to GRCh37, excluding translocations, and found 4,495 that fail. As these variants will increase our novelty estimate, we chose to exclude them for comparison with the published datasets. We did not include translocations in the comparison.

We used a threshold of 30% reciprocal overlap between variants identified in our dataset and either published callset to classify them as the same variant. This was implemented using bedmap v2.4.35 (Neph at el., 2012). For the comparison, we chose to be conservative by assessing whether a locus is structurally variable, rather than comparing the class of variant between the callsets. The reasoning for this is the possible misclassification of variant class (e.g. insertion vs duplication, in addition some inversions identified in the 1KG have since been shown to be inverted duplications and deletions (Soylev et al., 2019)). This analysis shows 78% of variants in our callset not to be present in either the 1000G or SGDP callsets. Some of these variants reach high frequency in regional groups or individual populations (Figure S9). To further evaluate the quality of our callset, we extracted common African variants in the 1KG that overlap common African-specific variants in our dataset, based on 75% reciprocal overlap (> 5% minor allele frequency). Although we expect some variation due to the different African populations in the two datasets, we should see a correlation at common variation. Indeed, we do find high correlation of allele frequencies between the two callsets (r = 0.72, Figure S3).

### Population Structure

We ran PCA using plink2 v2.00a2LM (Chang et al., 2015). We set variants with GQ < 20 to missing, included variants with minor allele frequency > 1%, missingness < 1% and pruned for linkage disequilibrium using the option --indep-pairwise 50 5 0.2. For the GenomeSTRiP dataset, we extracted biallelic deletions and biallelic duplications and ran the PCA separately. We excluded a single variant from the pruned duplication set due to it likely being affected by genotyping error (Hardy-Weinberg equilibrium (HWE) test = 1.73e-24). For deletions, we see clear patterns of structure across 10 principal components (Figure S4A). For multiallelic variants, we used a newer version of plink2 (v2.00a3LM) which can run PCA for multiallelic variants, and using the same parameters above.

Due to the relatively large number of PCs with observed patterns of structure, reflecting the diversity of our dataset, we ran a Uniform Manifold Approximation and Projection (UMAP) on the top 10 PCs that show population structure in the GenomeSTRiP deletion PCA (McInnes et al., 2018). This was implemented in R-3.6.0 using the package uwot (v0.1.3; https://cran.r-project.org/web/packages/uwot/index.html) setting initialization for the coordinates as ‘spca’, min_dist = 0.001, and n_neighbors = 16. As UMAP hyperparameters affect the local and global structure of the data, we present multiple figures with differing values for n_neighbors and min_dist (Figure S5).

For biallelic duplications, we see structure limited to the first four PCs (Figure S4B). However, the first two PCs separate Africans and Oceanians from the rest of the samples, in contrast to deletions. To further investigate this, we looked at the variant loadings in the PCA and find two variants with particularly high loadings, which when excluded, returns a similar, albeit much less defined, pattern to deletions. Those two variants were found to be the Oceanic-specific duplication on chr16p12 putatively introgressed from Denisovans and the highly differentiated *TNFRSF10D* variant. The relatively small number of large biallelic duplications identified renders the PCA sensitive to the few highly stratified variants found in Oceanians. A UMAP was run on the top 4 PCs as described above. The multiallelic variant UMAP was run on the top 10 PCs.

We also ran a PCA using the same parameters above on all classes identified in the Manta+Graphtyper callset (Figure S4C-E). We additionally excluded variants with HWE < 0.0001, and similarly to the GenomeSTRiP callset, we see population structure across all classes. However, we find more defined structure in the deletion Manta+Graphtyper callset in comparison to the GenomeSTRiP callset, which is likely due to the larger number variants identified by Manta. We ran a UMAP on the top 20 PCs deletion genotypes as implemented above and, as expected, see a more defined pattern of structure (Figure S6). We also ran a UMAP for insertions (top 10 PCs) using the same parameters. In Figure 1 of main text, the deletion and insertion UMAP was constructed from the Manta dataset, while the biallelic duplication and multiallelic variant UMAP was run using the GenomeSTRiP callset.

### Regional and Population-Specific Variation

We explored the total number and frequency of variants that are specific to continental and geographic regions (Figure S8). As this analysis is sensitive to individuals with recent admixture, we used previous estimated individual ancestry from SNV analysis and excluded samples that show such admixture (for more details refer to Bergström et al., 2019). To further conservatively avoid over-counting single variants that have been called as multiple adjacent entries, potentially as a result of a complex structural event, we merged variants with similar allele frequencies and the same copy number lying within 25 kb of each other.

For multiallelic copy number variants, we restricted the analysis to high quality variants that have CNQ ≥ 13. This score is phred-scaled, with CNQ ∼ 13 representing ∼95% confidence of diploid copy number. In the expansion plots presented (Figure 3 and Figure S11), the highlighted regions (red bar) illustrate the expanded regions. However, in some cases the discovery algorithm finds the expanded region to vary in size and can be slightly larger in different samples. To be conservative we display the smallest overlapping region consistent across samples.

Similarly, we explored the total number and frequency of variants that are only found in a specific population (Figure S7). Here, we did not exclude samples based on known admixture as in the regional analysis. We note that this analysis is sensitive to the sampling location and sample size of each population, i.e. if a region is more comprehensively sampled we would expect a lower number of population-private variants in contrast to more sparsely sampled regions. In addition, even if we find a variant that seems population specific, it may be present at a lower frequency in another population but was not captured due to sample size. Nevertheless, we still identify examples of high-frequency population-specific variants that are not found in geographically nearby populations.

#### *OCA2* deletion in BantuSouthAfrica

We find a 2.7 kb deletion in *OCA2* (also known as *P* gene) to be of surprising frequency (44%) and an outlier exclusive in the Bantu South African population (Figure S7). This deletion has been reported previously in African populations, and is known to cause Brown Oculocutaneous Albinism following a recessive mode of inheritance (Durham-Pierre et al., 1994; Manga et al., 2001). We find homozygotes for this deletion in our dataset, suggesting that samples with albinism were donated to the HGDP collection. We contacted CEPH (Centre d’Etude du Polymorphism) about this observation and were informed that Trefor Jenkins (now deceased) was the researcher who provided samples from this population. As he has a history of working with African populations with albinism (Stevens et al., 1997), we conclude that this variant in the HGDP dataset is likely to result from the particular sample ascertainment rather than being representative of its frequency in the Bantu South African population.

#### *SIGLEC5* deletion in Mbuti

In the main text, we report a deletion that is specific and high frequency (54%) in the Mbuti population that deletes the inhibiting receptor *SIGLEC5* without removing its paired activating receptor *SIGLEC14* (Ali et al., 2014). We also find a previously reported deletion which removes the function of the activating Siglec-14, to be common in all populations (global frequency 38%), with particularly high frequency in East Asians (63%). This common deletion removes the activating receptor, by fusing *SIGLEC5* and *SIGLEC14*, creating a gene that has the *SIGLEC5* coding sequence and expressed under the promoter of *SIGLEC14* (Yamanaka et al., 2009). We discover a single Mbuti sample (HGDP00450) that has both deletions on separate haplotypes. By looking at depth in this region the two deletions appear complex, but we were able to resolve them using 10x linked reads (Figure S14).

### Population Stratification

We calculated the maximal allele frequency difference for each population pair (total 1431 pairwise comparisons) and assessed this in relation to the average SNV differentiation between each population (SNV Fst). SNV Fst was calculated using EIGENSTRAT on all SNPs within the accessibility mask defined in Bergström et al. 2019 (Price et al., 2006). We calculated structural variant allele frequency and missingness in each population separately setting variants with GQ < 20 to missing, and excluded variants with missingness > 25% in each population. We then calculated the maximal variant allele frequency difference for each population pair, separately for deletions (which include biallelic deletions and deletions in multiallelic sites), biallelic duplications and insertions. As these values are sensitive to the sample size of each population, we assessed this relationship in Figure S10. For the *HBA2* deletion we find almost fixed in the PapuanSepik population, we find the PapuanHighlanders, who do not have the deletion, have high missingness at this variant. The variant GQ is 19 for almost all samples in this population, just missing the threshold we set. However, closer inspection of the variant quality shows that the copy number genotype quality (CNQ) was high for these individuals (all CNQ > 70, except one CNQ = 18). Thus this variant was subsequently included in this population for analysis.

### Archaic Introgression

We genotyped the identified CNVs from this study (GenomeSTRiP calls) in the three published high coverage archaic genomes: Altai Denisova (Meyer et al., 2012), Altai Neanderthal (Prufer et al., 2014) and Vindija Neanderthal (Prufer et al., 2017). In these previous studies, sequencing reads for each sample were aligned to GRCh37. For our analyses, we downloaded these mapped reads and remapped them to GRCh38 using bwa aln v0.7.12 (Li & Durbin 2009), with parameters tuned for ancient DNA (“-l 16500 -n 0.01 -o 2”), and marked duplicates using the MarkDuplicates tool from Picard v2.6.0 (http://broadinstitute.github.io/picard/). Each ancient genome was joint-called separately using a site VCF with 30 Sanger-PCR samples using GenomeSTRiP. This was done as we were concerned that the different library preparations of the HGDP dataset may affect calling in the archaic genomes, so we limited joint calling to single library (Sanger-PCR) and a single archaic genome. We then investigated variants that were highly stratified (Vst > 0.2) and shared with any archaic genome but missing from African populations, and restricted analysis to high quality archaic variant calls (CNQ ≥ 13). All identified putative introgressed variants were then checked and confirmed manually in IGV (Thorvaldsdóttir et al., 2013). In the Manta dataset we identified a relatively small deletion within *JAK1* (375 bp) which is specific to Oceania at 44% frequency. We checked the archaic genomes in IGV and find the Denisovan genome to be homozygous for the deletion, and the Altai Neanderthal to be homozygous reference. The Vindija Neanderthal shows a less clear genotype: we do see a reduction in depth relative to flanking regions; however, due to the small size of the region it is difficult to ascertain if it is heterozygous for the deletion or if the reduction in depth is due to stochastic noise. To be conservative we do not consider the Vindija Neanderthal genotype in Table 1 in the main text. We also identify small deletion (63 bp) within *ZNRF1* specific to Oceania at 34% frequency. Manually checking the variant using IGV in the archaic genomes illustrated that all three are homozygous for the deletion. We present region screenshots showing examples of variants identified in modern and archaic genomes (Figure S15-17.)

For the chr16p12 duplications exclusive to Oceanians, we used the estimated individual ancestry from SNV analysis (Bergström et al., 2019) and find that all 11 Bougainville samples have appreciable East Asian ancestry (average 19%, minimum 16%, maximum 21%), one out of the eight Sepik/Lowlanders had 20%, while all eight Highlanders show no East Asian component.

### Longranger and Supernova Assembly

In the SNV analysis (Bergström et al., 2019), 26 HGDP samples from 13 populations (two per population) were sequenced using 10x Genomics linked-reads. For the present study, we performed an additional lane of sequencing for these 26 samples from the same library preparation to increase coverage for structural variants analysis and additionally for the *de novo* assembly using the Supernova assembler v2.1.1. We used the Long Ranger v2.12 pipeline which generated phased VCFs of structural variants. This was performed twice, once for single-lane and another for two-lane (higher coverage) libraries. In this study, we use the linked reads to validate variants we identified in the standard Illumina WGS and present it as a resource for the scientific community. We also used linked reads from two lanes as input to Supernova v2.1.1 and selected the pseudohap2 output which extracts both pseudohaplotypes (Weisenfeld et al., 2017). Assembly statistics are presented in Table S2. We observe variable contiguity and assembly sizes for the assemblies, likely reflecting the initial average molecular size for each sample. One sample had a markedly smaller assembly size compared to the rest (HGDP00954), and was excluded for assembly based analysis, leaving 25 samples.

### Non-Reference Unique Insertions

To identify non-reference unique (non-repetitive) insertions (NUIs), we used the NUI pipeline which compared each of the Supernova assemblies to the GRCh38.p12 reference (Wong et al., 2018, https://github.com/wongkarenhy/NUI_pipeline). We followed the definition of NUI as proposed by Wong et al., 2018. Sequencing reads were extracted from BAM files generated from samples sequenced using one lane by the Long Ranger v2.12 pipeline. Briefly, the pipeline takes poorly mapped, unmapped and discordant reads from the Longranger output and maps them to the Supernova assemblies. It subsequently identifies read clusters and extends the contig ends to use as anchors, which are then aligned against GRCh38. Breakpoints are subsequently identified and filtered to identify NUIs. The NUIs are then blasted to GRCh38p.12, including all the patches, to confirm they are not present in the reference. The number of NUIs per sample is shown in Table S2. To assess the potential functional effect of each identified insertion we used the Variant Effect Predictor (McLaren et al., 2016). We set the “Upstream/Downstream distance (bp)” = 0 and extracted “canonical” transcripts from the predicted results. To identify if coding sequences are affected, we filtered for “coding_sequence_variant”. Some insertions affected more than one transcript. For PCA we excluded variants that are present in four or less individuals or that are present in more than 23 individuals and used the prcomp function in R-3.6.0 with default parameters. We show PC3-4 as the top two PCs likely represent variation in assembly size and quality, with correlation observed between PC2 values and contig N50 (r = 0.63). Additionally, the number of identified insertions is correlated with the contig N50 (r = 0.91, Figure S18). NUIs density across chromosomes was plotted using karyoploteR v1.10.4 (Bernat and Serra, 2017).

### Fluorescent in situ hybridisation (FISH)

Melanesian lymphoblastoid cell lines were purchased from Coriell Institute for Medical Research (GM10543 and GM10540) while fosmid and bacterial artificial chromosome (BAC) clones used in this study were provided by the clone archive team of the Wellcome Sanger Institute (Table S3). Fosmid/BAC DNA was prepared using the Phase-Prep BAC DNA kit (Sigma-Aldrich) following the manufacturer’s protocol. For fibre-FISH, stretched chromatin and DNA fibres were prepared by alkaline lysis of lymphoblastoid cells deposited on Thermo Scientific™ Polysine adhesion slides (Fisher Scientific) as described previously (Korbel et al., 2007). Purified fosmid/BAC DNA were first amplified using the GenomePlex® Complete Whole Genome Amplification kit (WGA2) (Sigma-Aldrich) and then labelled with either biotin-16-dUTP, Dinitrophenol (DNP)-11-dUTP or Digoxigenin (DIG)-11-dUTP (Jena Bioscience) using the GenomePlex® Complete Whole Genome Reamplification kit (WGA3) (Sigma-Aldrich) as described in Louzada et al., (2017). The DNP-labelled probes were detected with rabbit anti-DNP and Alexa 488 conjugated goat anti-rabbit IgG (Invitrogen). The DIG-labelled probes were detected with monoclonal mouse anti-DIG IgG (Sigma-Aldrich) and Texas red conjugated donkey anti-mouse IgG (Invitrogen). The biotin-labelled probes were labelled with biotin-16-dUTP and detected with one layer of Cy3-streptavidin (Sigma-Aldrich). After detection, slides were mounted with SlowFade Gold® (Invitrogen) mounting solution containing 4′, 6-diamidino-2-phenylindole (Invitrogen). Metaphase chromosomes were prepared from lymphoblastoid cell lines following standard procedure (Howe et al. 2014). Metaphase- and interphase-FISH essentially followed Gribble et al., (2011). Probes directly labelled ChromaTide™ Texas Red®-12-dUTP (Invitrogen), Green-dUTP (Abbott), Cy3-dUTP and Cy5-dUTP (Enzo) were used in this study. Images were captured on a Zeiss AxioImager D1 fluorescent microscope and processed with the SmartCapture software (Digital Scientific UK).

---

**Figure S1:**
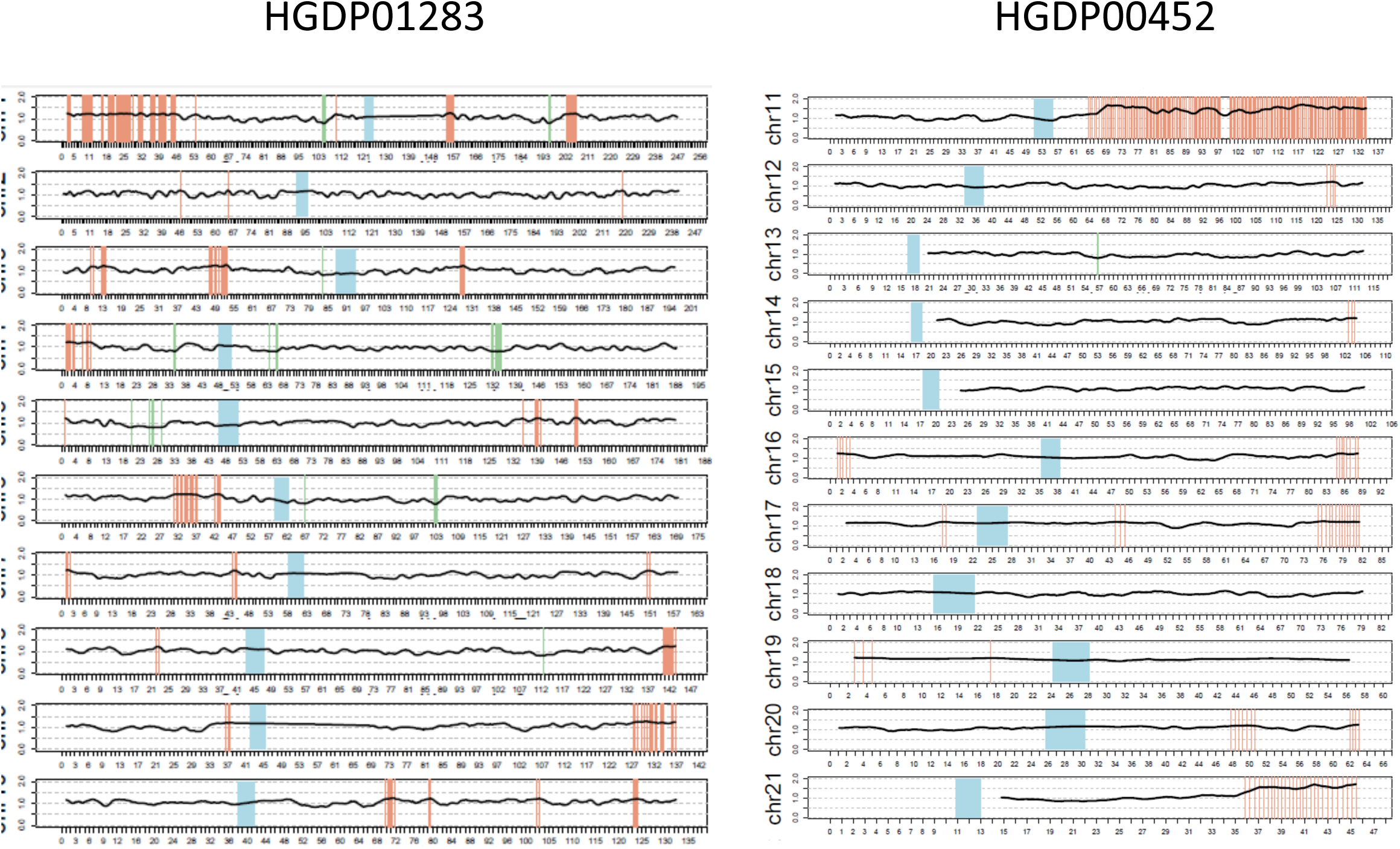
Coverage plots illustrating examples of samples that were excluded from the analysis because of likely cell-line artefacts. Coverage was calculated at ∼300,000 single positions across the genome and a rolling mean was plotted normalized by the genome-wide median. **Left**: HGDP01283 (chr1-10) which shows artefacts across multiple chromosomes. **Right**: HGDP00452 (chr11-21) which shows a large duplication in most of chromosome 11 in addition to smaller duplications in other chromosomes. Orange bars indicate coverage is >25% than chromosome average, green <25%. Blue represents centromeres.

**Figure S2:**
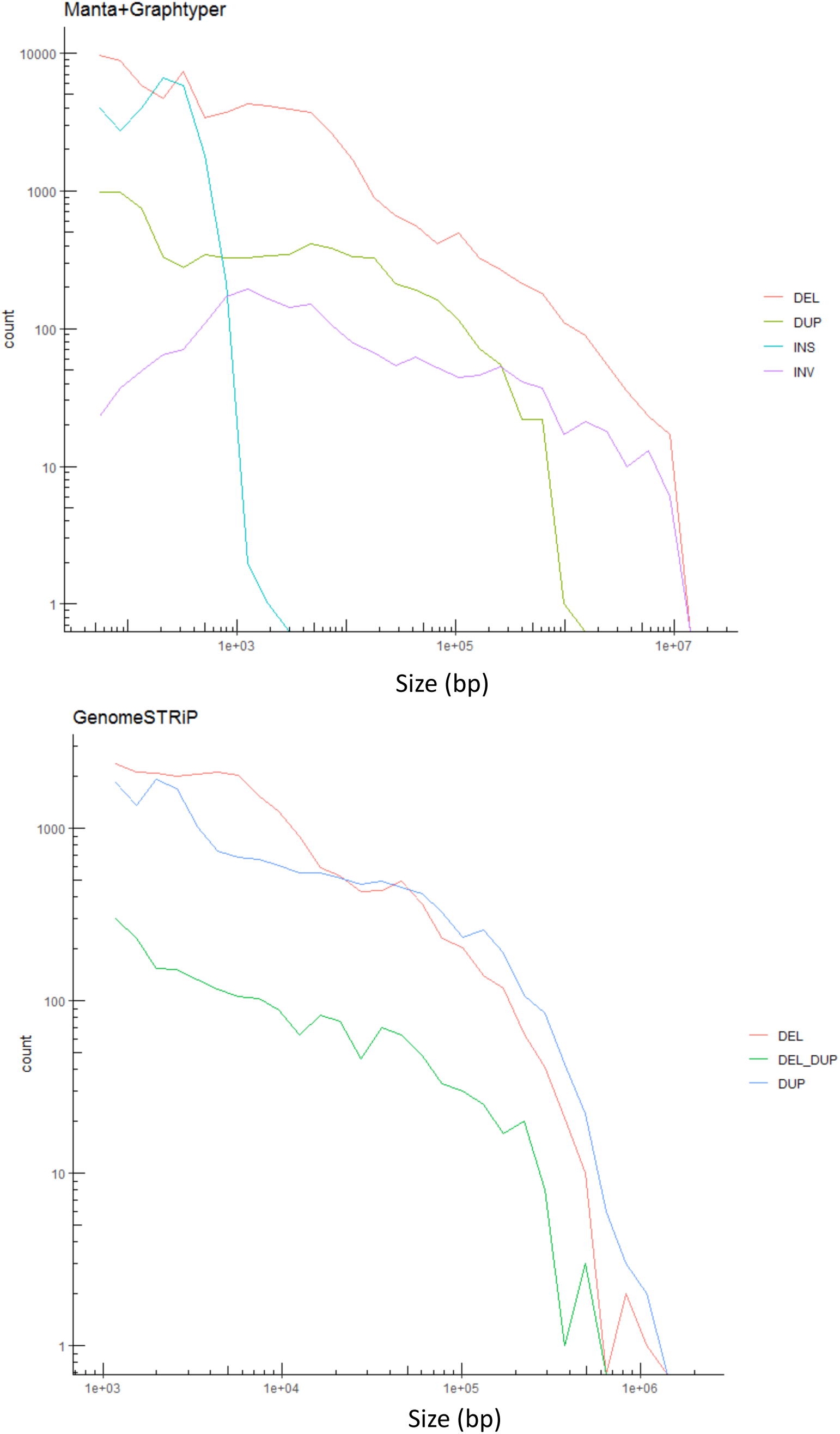
Size distribution of identified variants that passed all filters and were included in the final callset. Note the differences in scales between the two plots. **Top**: Manta+Graphtyper. **Bottom**: GenomeSTRiP – green line shows variants that have both deletion and duplication alleles.

**Figure S3:**
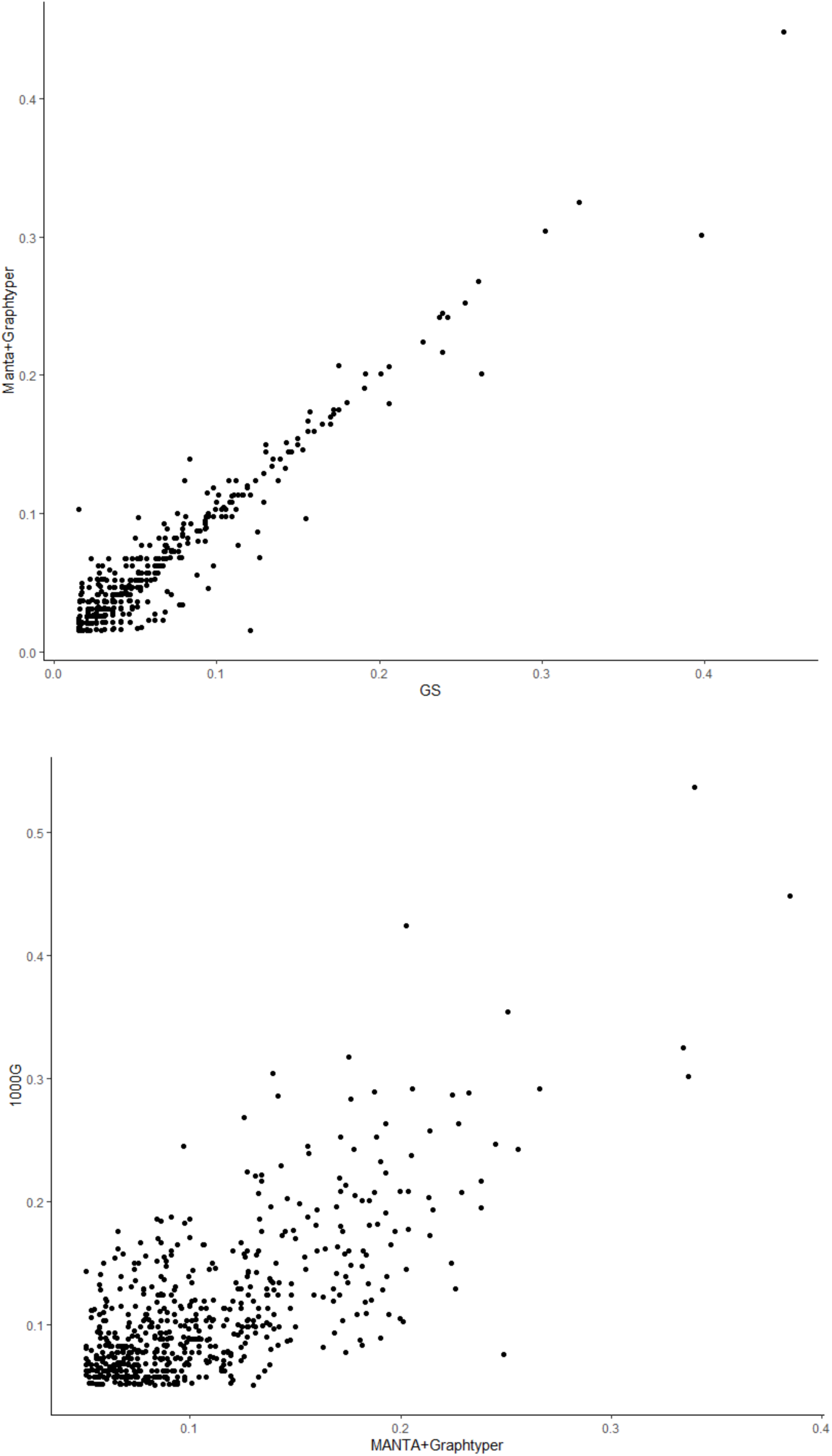
Quality checks on genotyping. **Top**: Correlation of allele frequency of variants identified by both Manta+Graphtyper and GenomeSTRiP (African-specific variants). **Bottom**: Allele frequency correlations between variants identified in the 1000G and the HGDP Manta+Graphtyper callset (common African-specific variants).

**Figure S4A:**
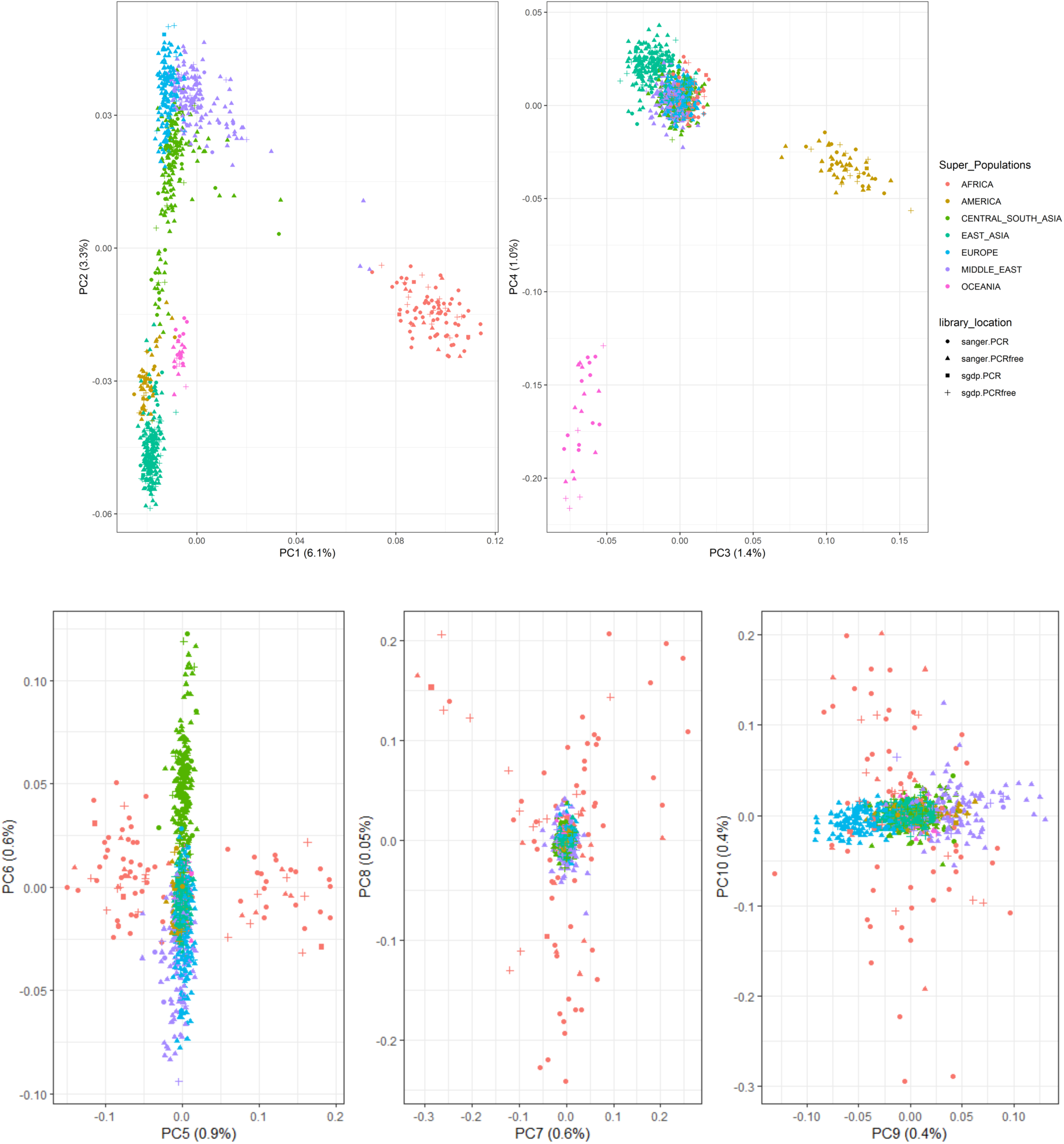
PCA (1-10) of GenomeSTRiP biallelic deletion genotypes by sample library preparation and sequencing location.

**Figure S4B:**
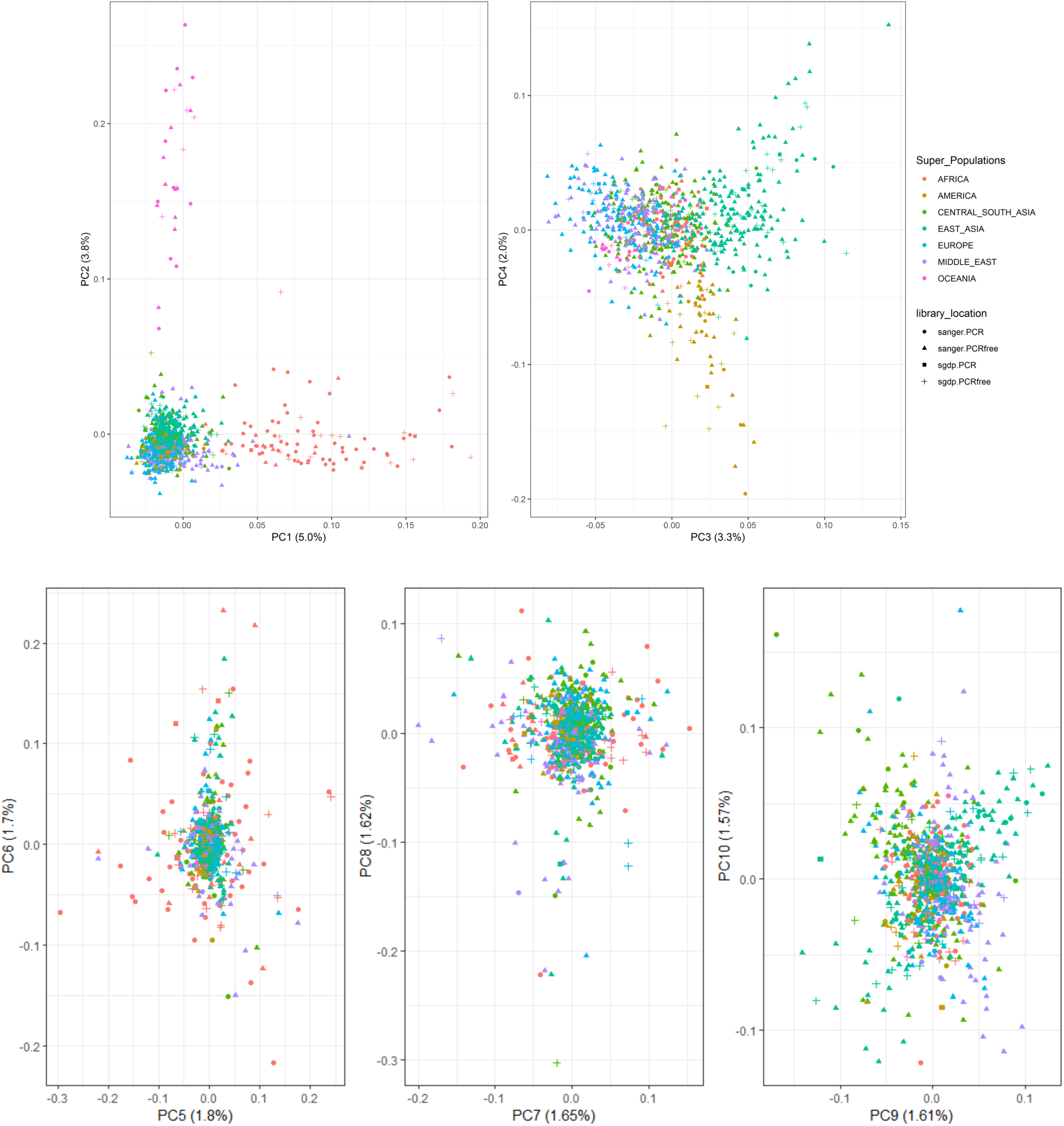
PCA1-10 of GenomeSTRiP biallelic duplication genotypes by sample library preparation and sequencing location.

**Figure 4C:**
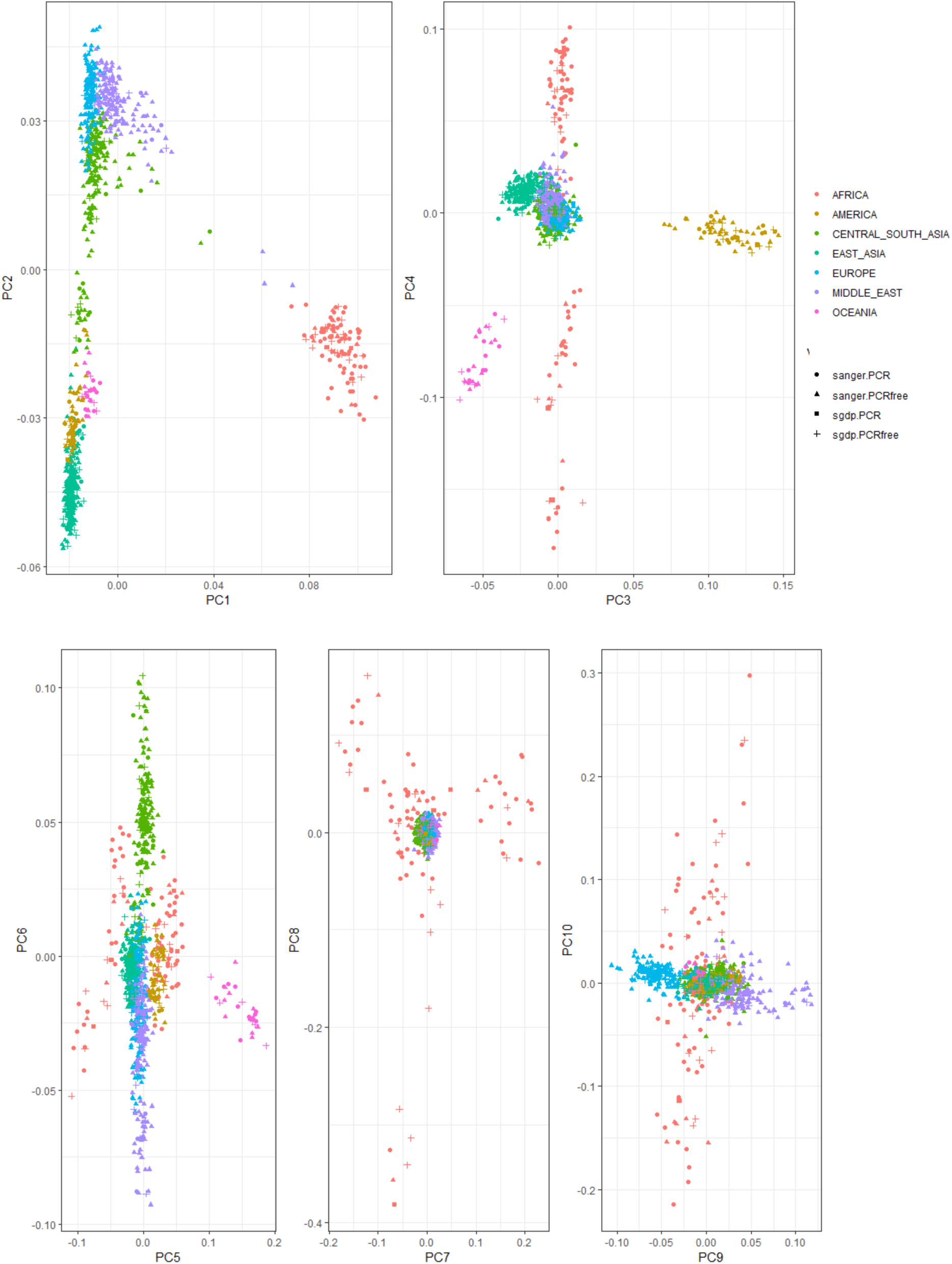
PCA1-10 of Manta+Graphtyper deletion genotypes by sample library preparation and sequencing location.

**Figure S4D:**
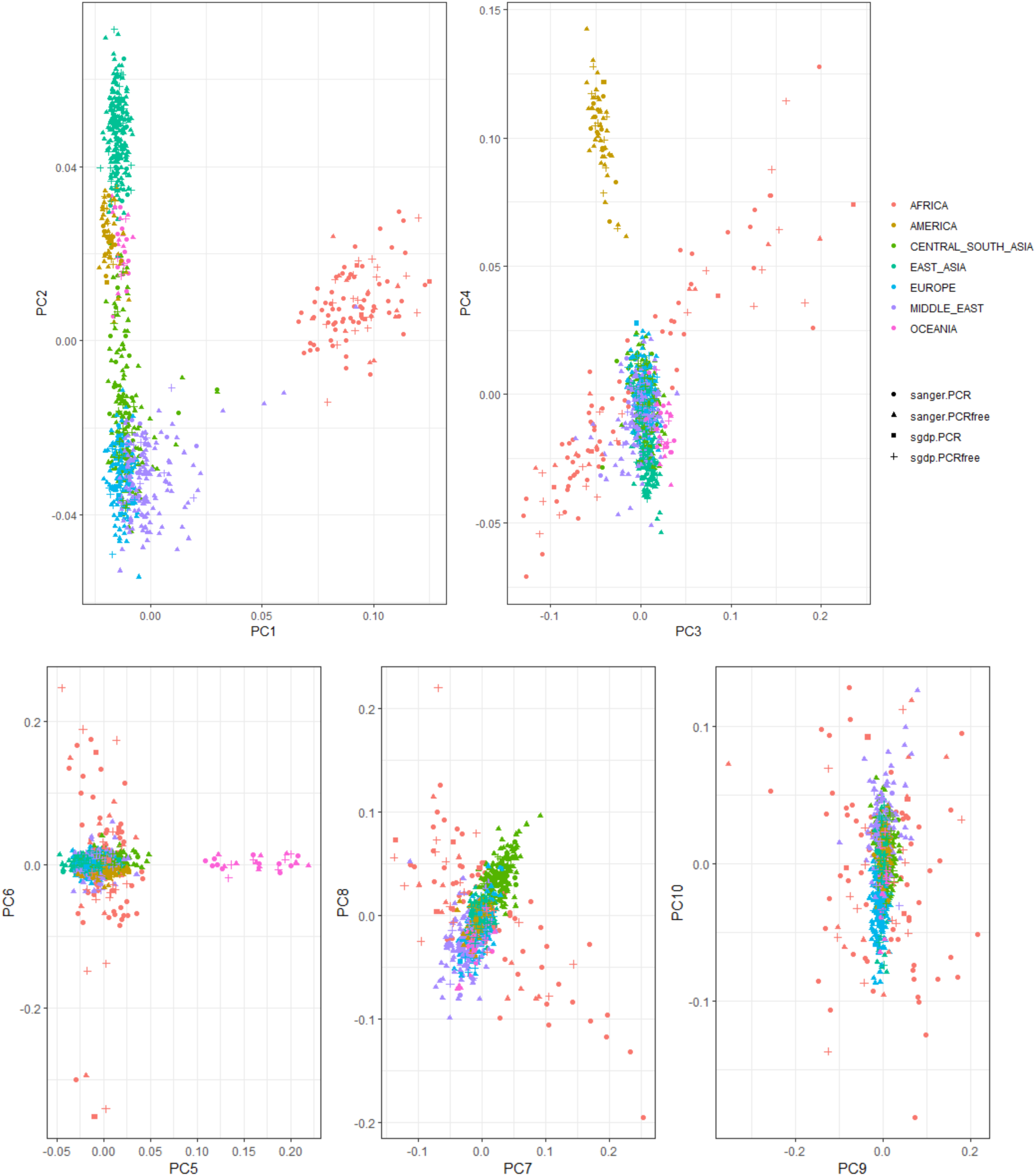
PCA1-10 of Manta+Graphtyper insertion genotypes by sample library preparation and sequencing location.

**Figure S4E:**
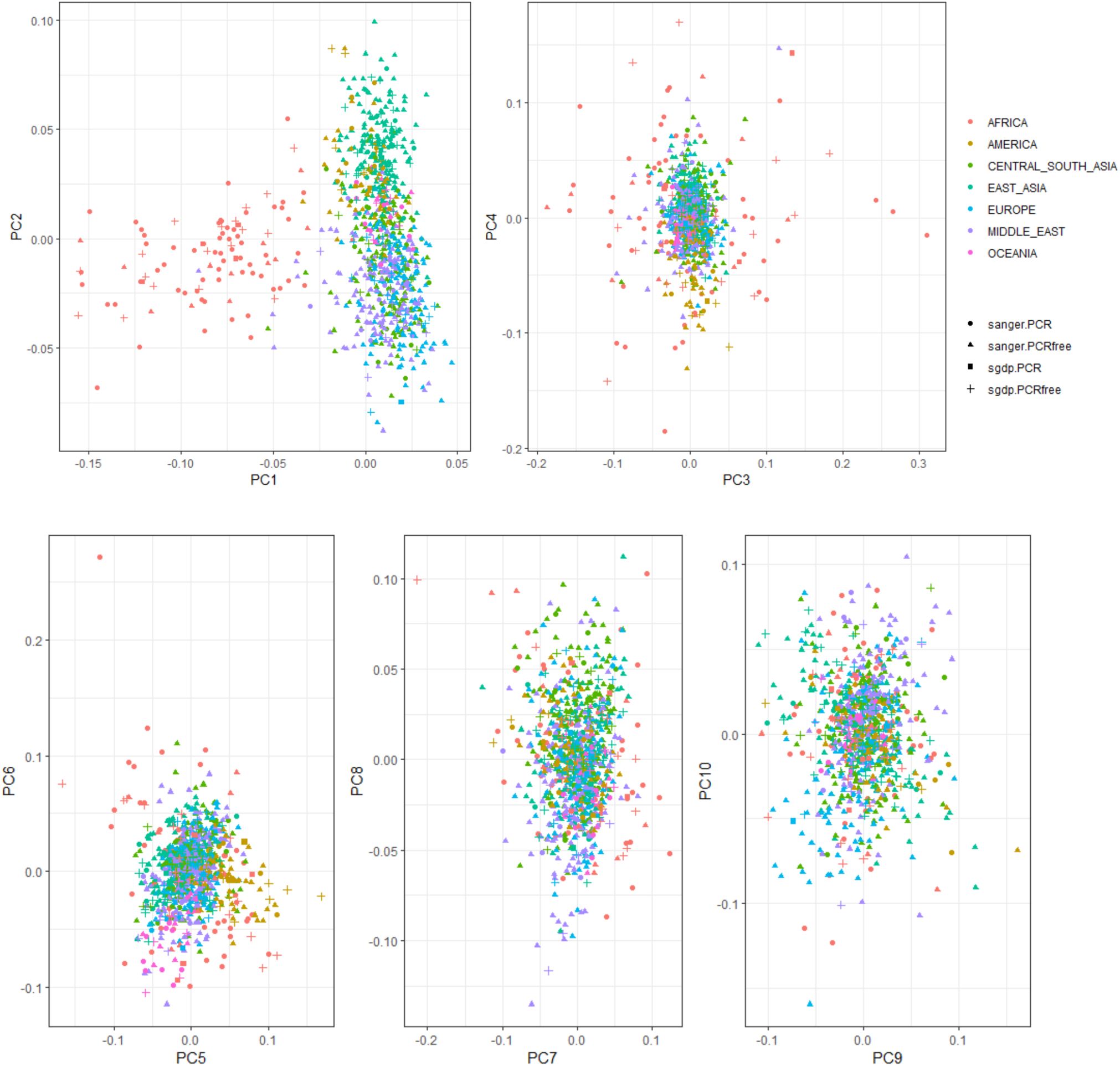
PCA1-10 of Manta+Graphtyper inversion genotypes by sample library preparation and sequencing location.

**Figure S5:**
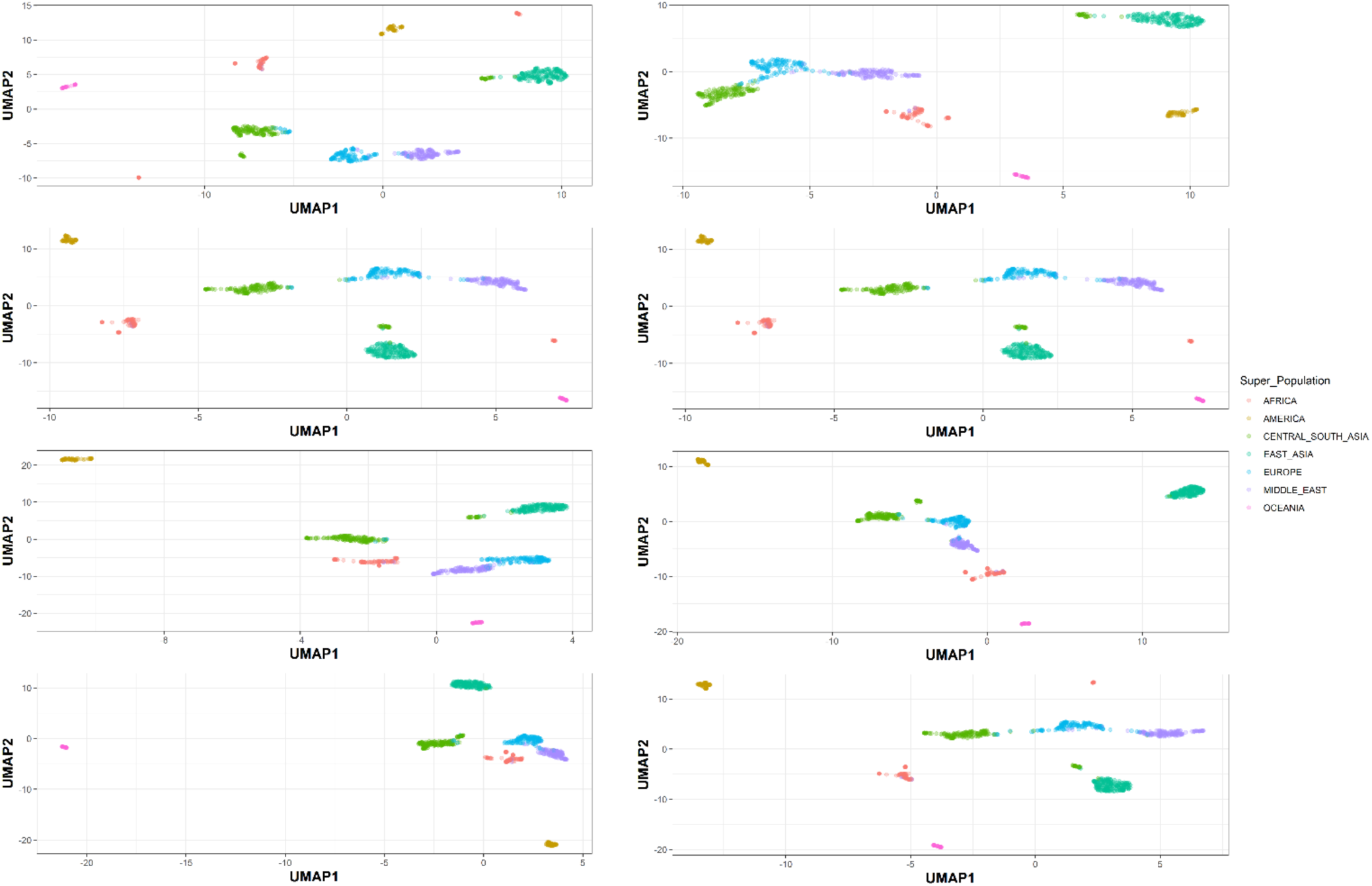
Effect of UMAP hyperparameters on observed clustering – GenomeSTRiP deletions. Left: Different n_neighbors values (increasing from top to bottom: 8,16,32,64) – keeping min_dist = 0.01 Right: Different min_dist values (increasing from top to bottom : 0.1,0.01,0.001,0.0001) – keeping n_neighbors = 16.

**Figure S6.**
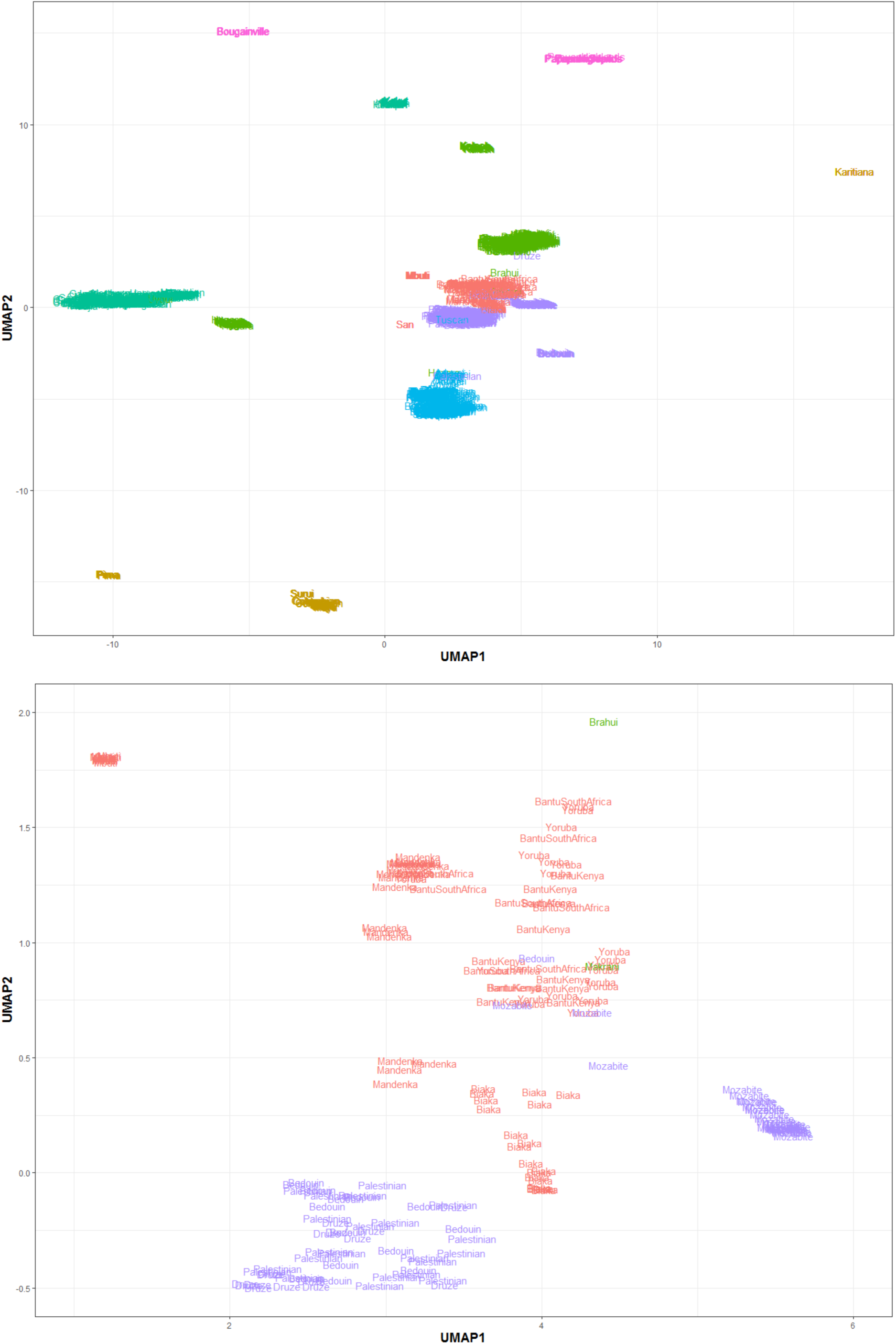
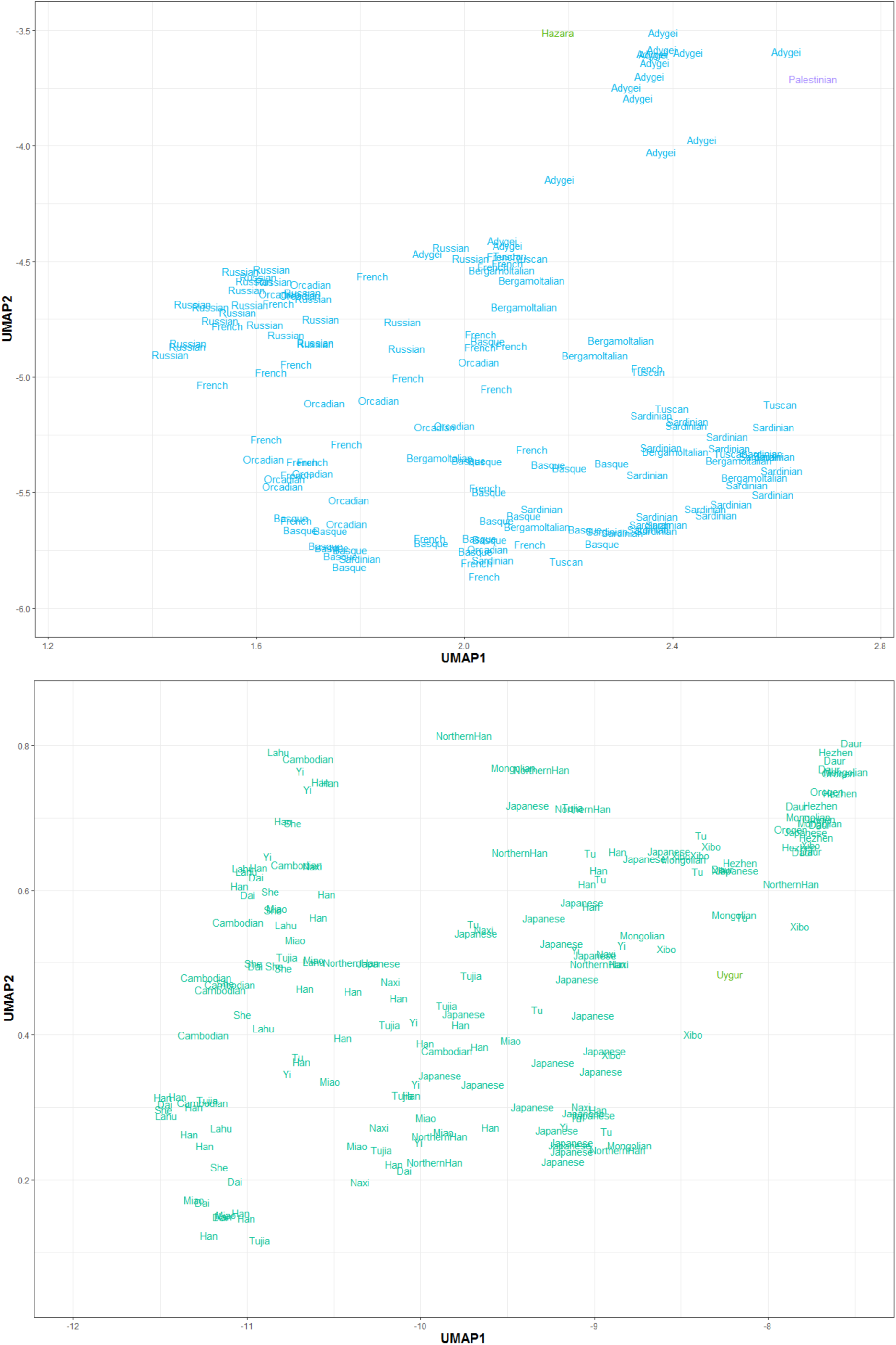
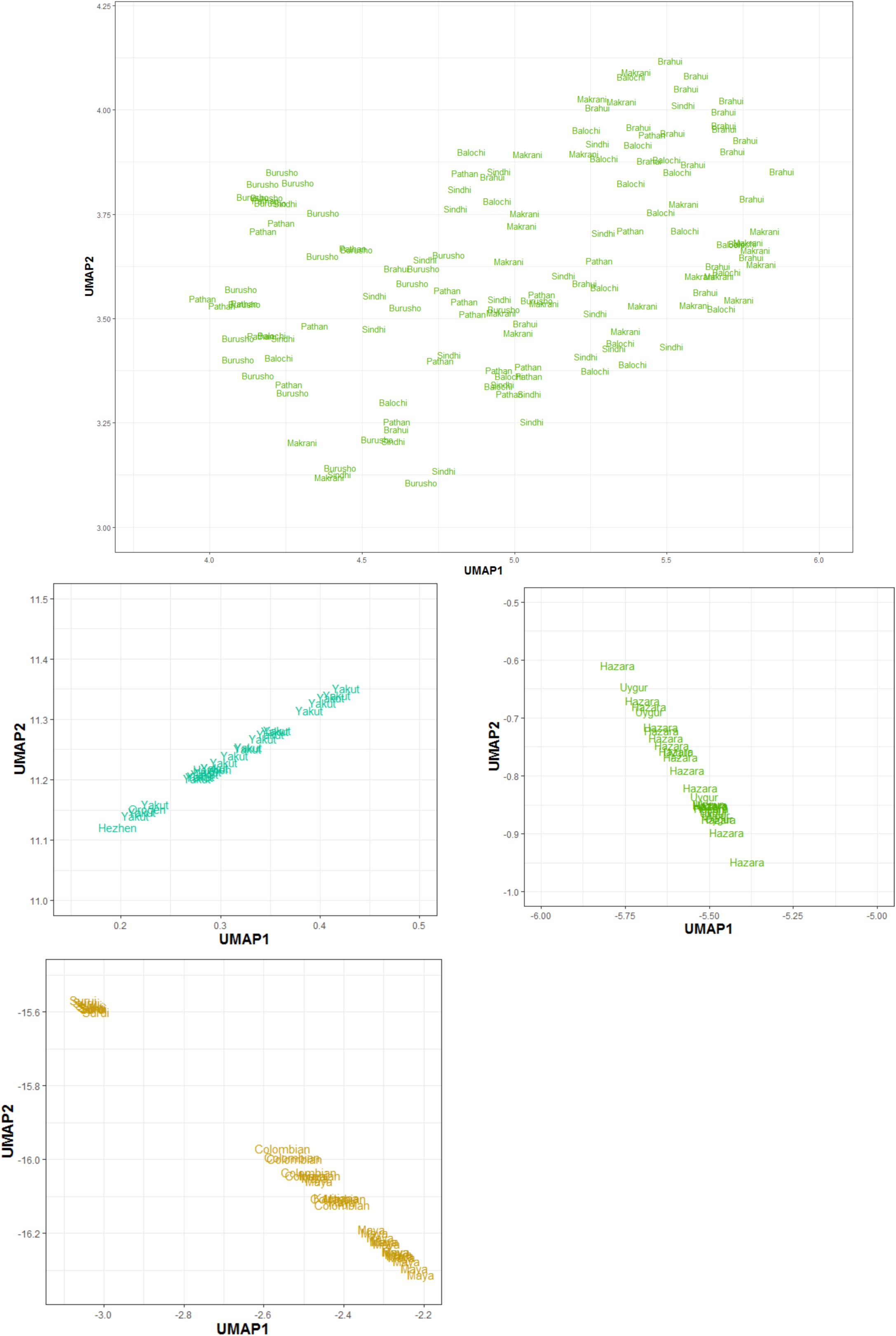
Fine structure in UMAP. Zoomed plots of each continental cluster with population labels from Figure 1 in main text. Based on Manta+Graphtyper deletion genotypes.

**Figure S7:**
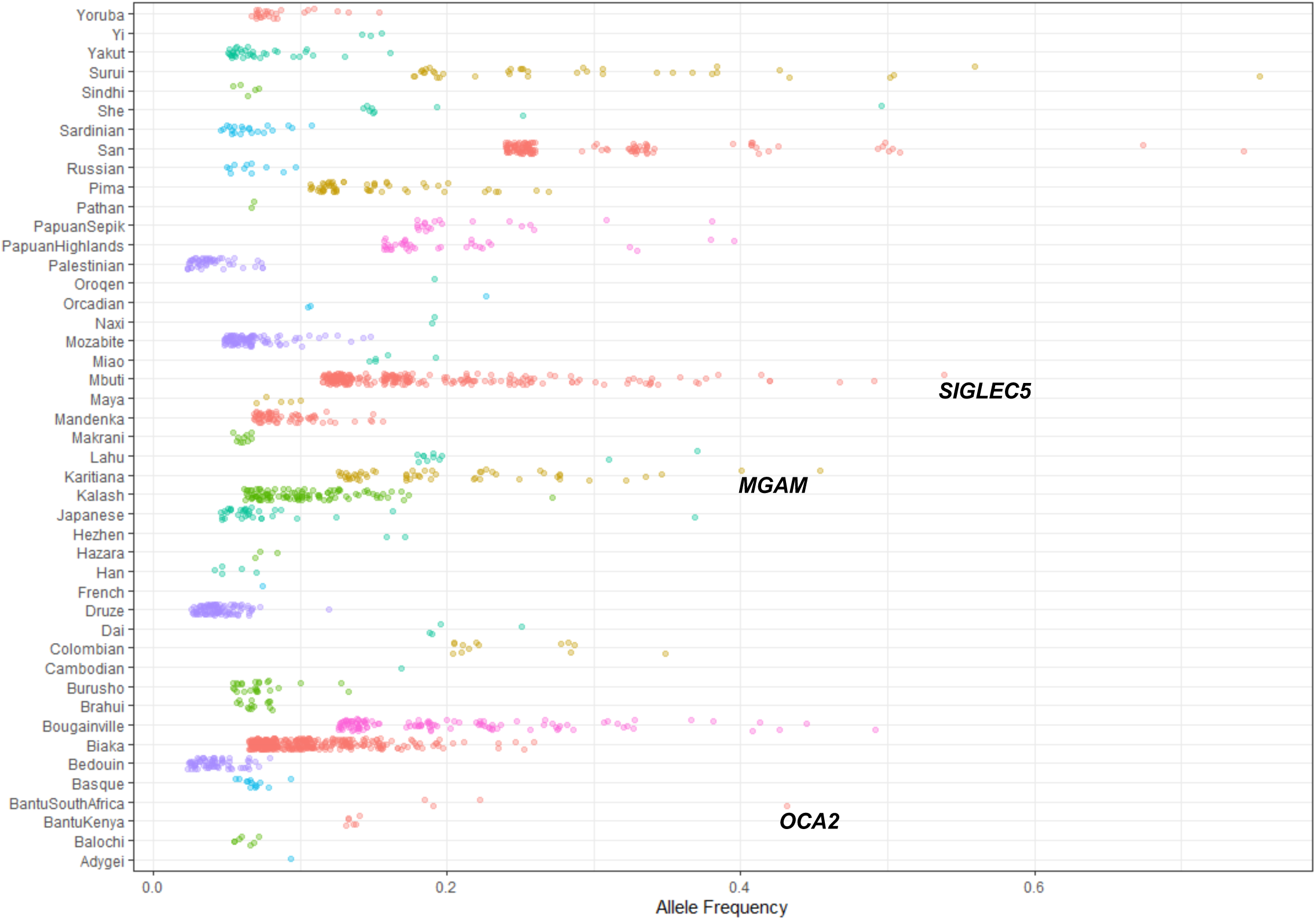
Population-Specific Variation – Each point represents a variant private to a population (n > 2) with the x-axis reflecting its frequency. Colours represent regional labels and random noise is added to aid visualization. High-frequency variants discussed in the text are highlighted.

**Figure S8:**
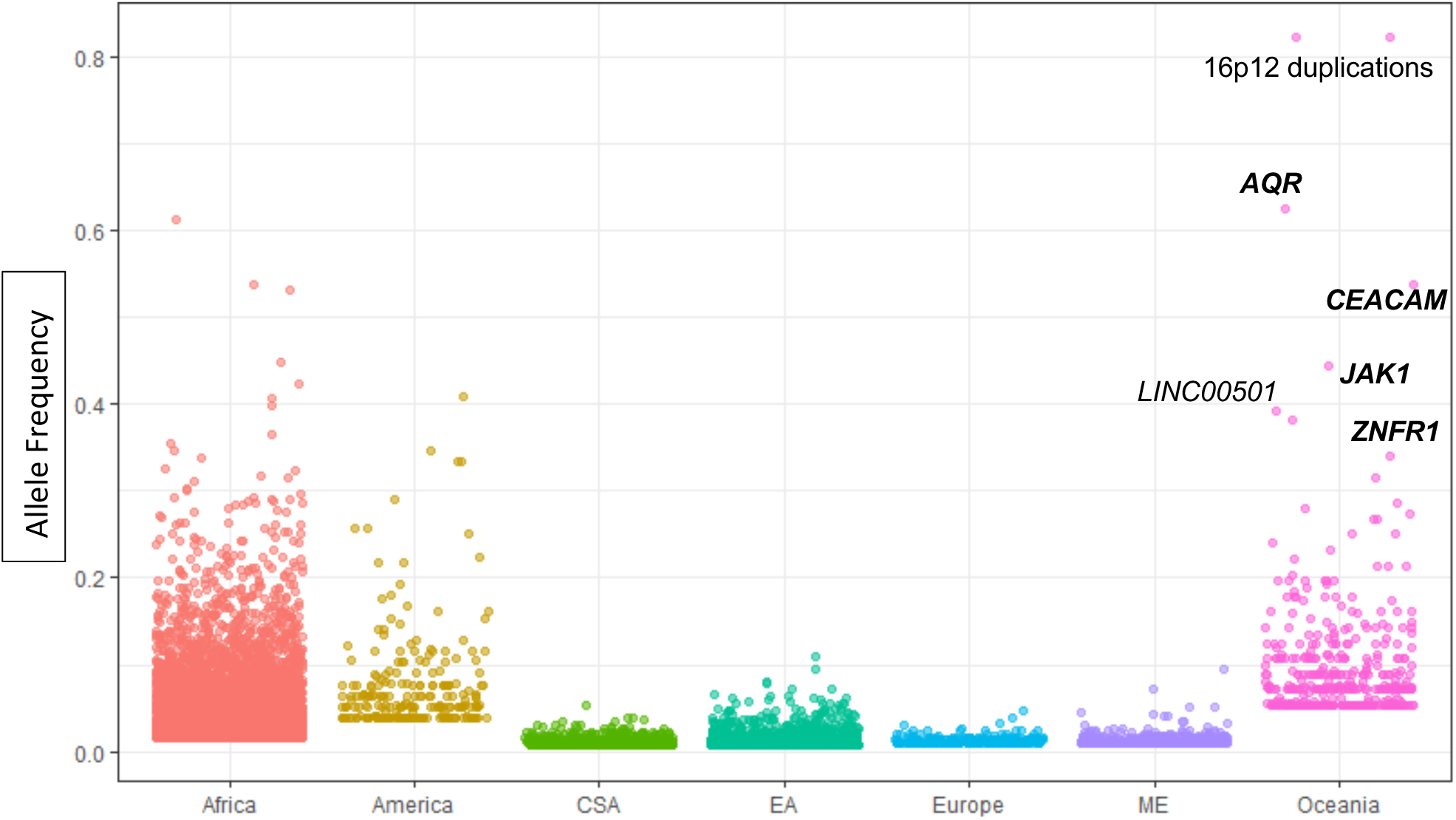
Regional-Specific Variation – Each point represents a variant private to a regional group (n > 2) with the y-axis illustrating its frequency. Random noise is added to aid visualization. The distribution reflects the ancestral diversity in Africa, the connectivity of Eurasia, the isolation & drift of the Americas and Oceania, and the separate Denisovan introgression event in Oceania. Oceania is notable for having private high-frequency variants that are all shared with the Denisovan genome and are within (**bold**) or near the illustrated genes, four of which are newly identified in this study (*AQR*, *CEACAM*, *JAK1, ZNFR1*). The Americas contain high frequency variants which are not shared with any archaic genomes, suggesting they arose and increased to high-frequency after they split from other populations. EA: East Asia, CSA: Central & South Asia, ME: Middle East.

**Figure S9:**
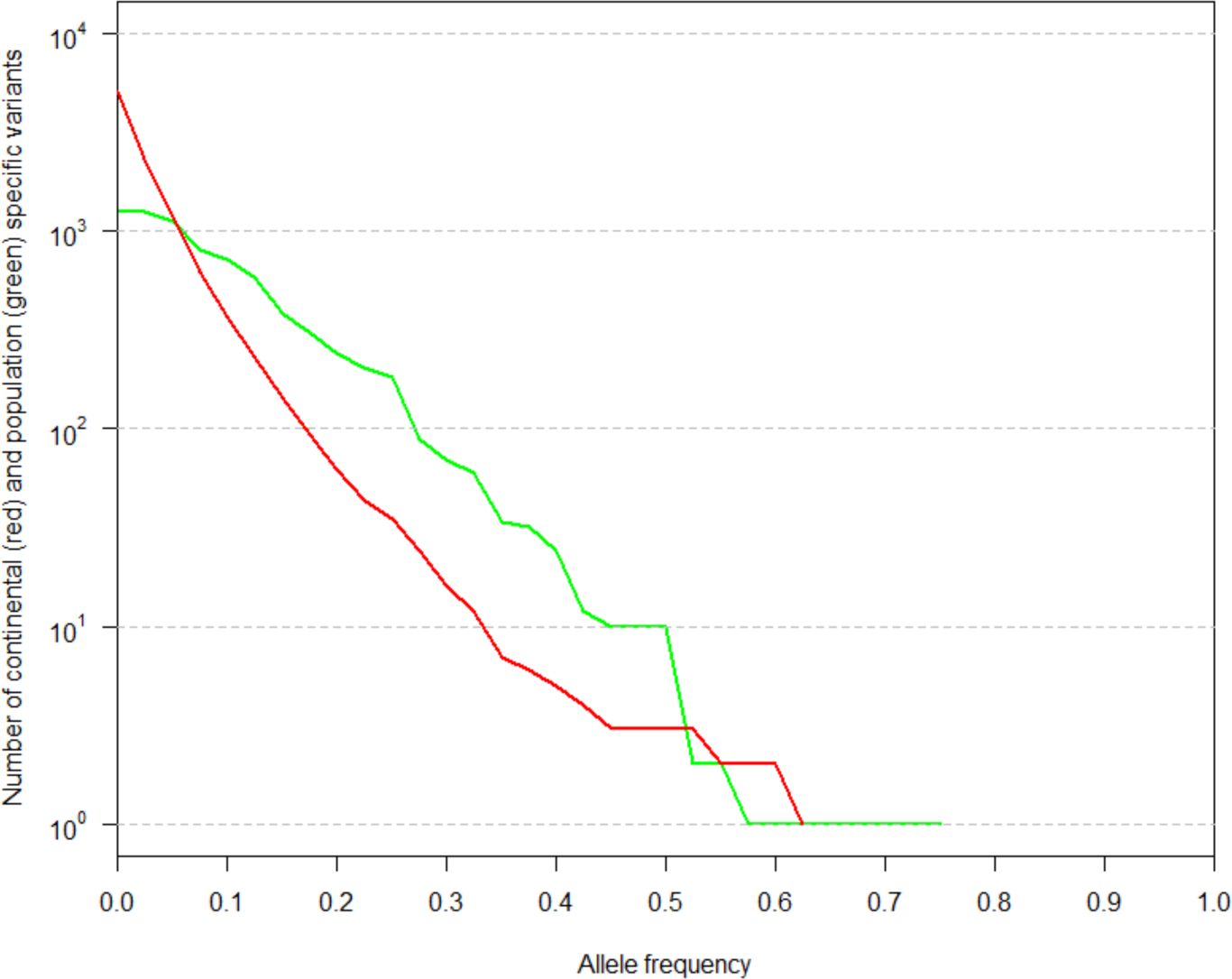
Continental (red) or Population (green) specific variants (n > 2) in the HGDP not found in 1000G or SGDP SV callsets binned by allele frequency. The same variant can be present in both distributions.

**Figure S10:**
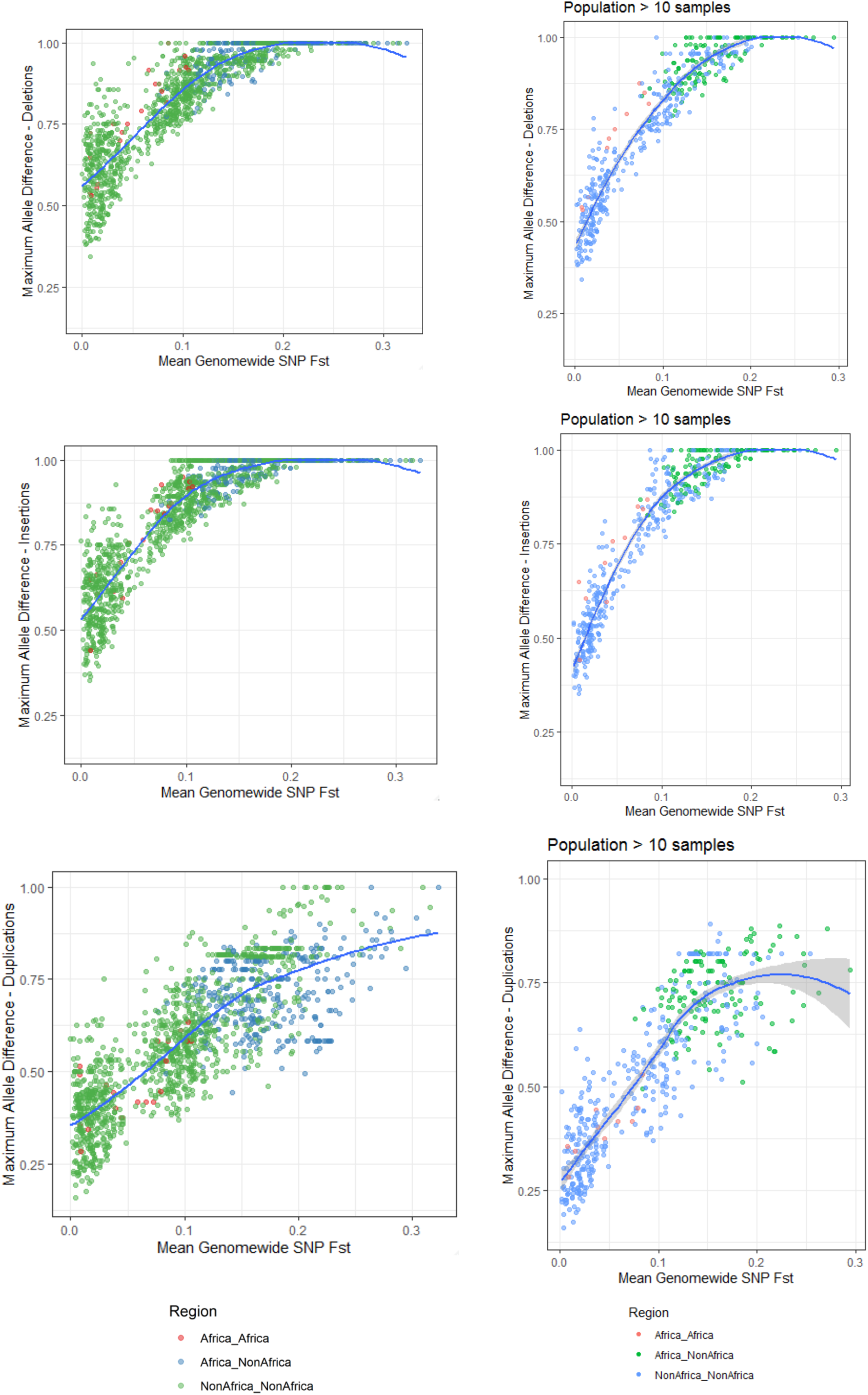
Maximum allele frequency difference as a function of population differentiation. Blue line is loess fits. **Left**: including all populations – **Right**: after excluding populations with 10 samples or less. Deletions (Top), Insertions (Centre), Duplications (Right).

**Figure S11:**
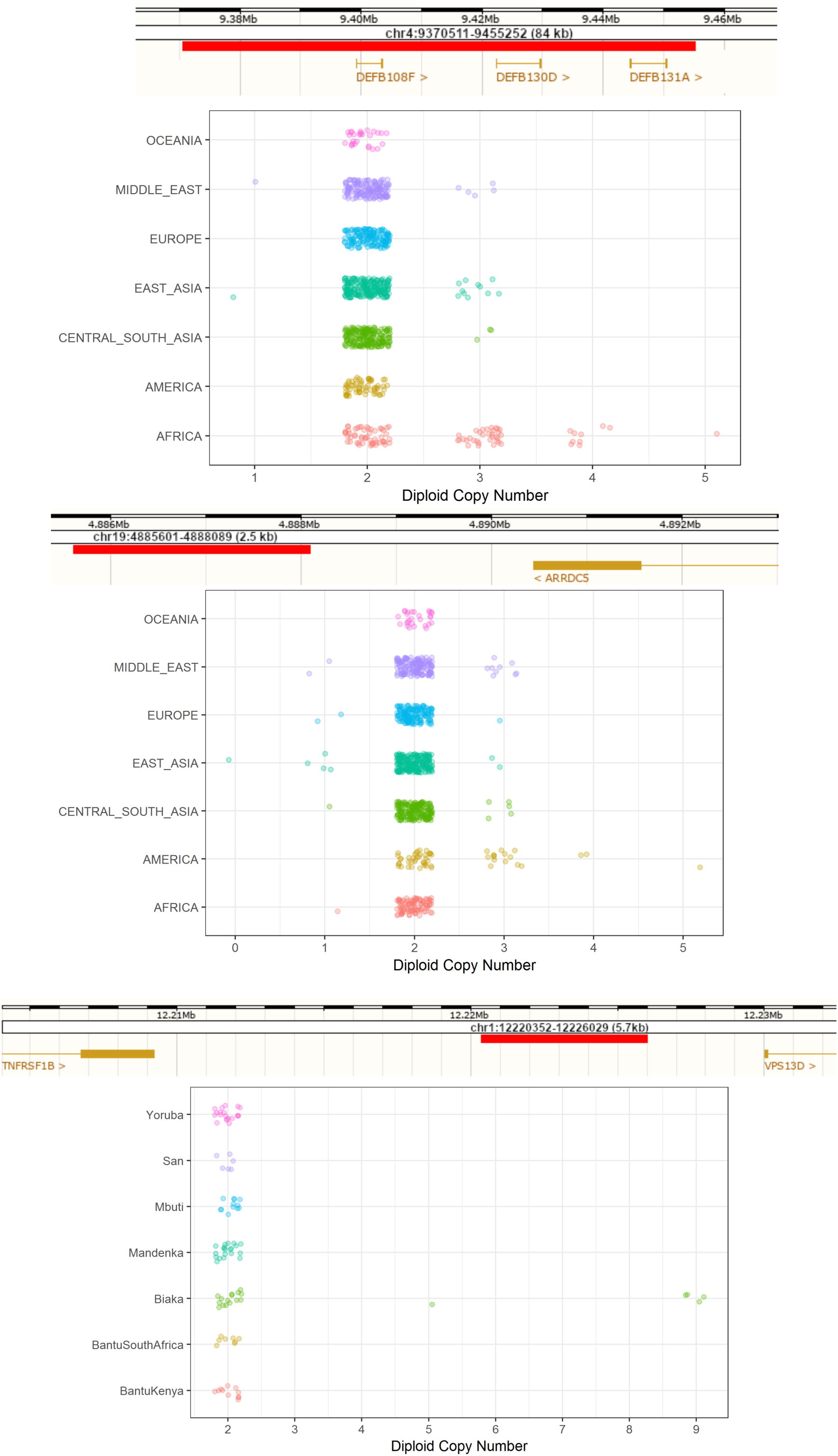
Additional copy number expansions. Red bar illustrates region expanded. **Top:** Expansions in beta-Defensin genes. **Centre:** Expansions downstream of *ARRDC5* prominent in Americans. **Bottom:** Expansion downstream *TNFRSF1B* private to Biaka.

**Figure S12:**
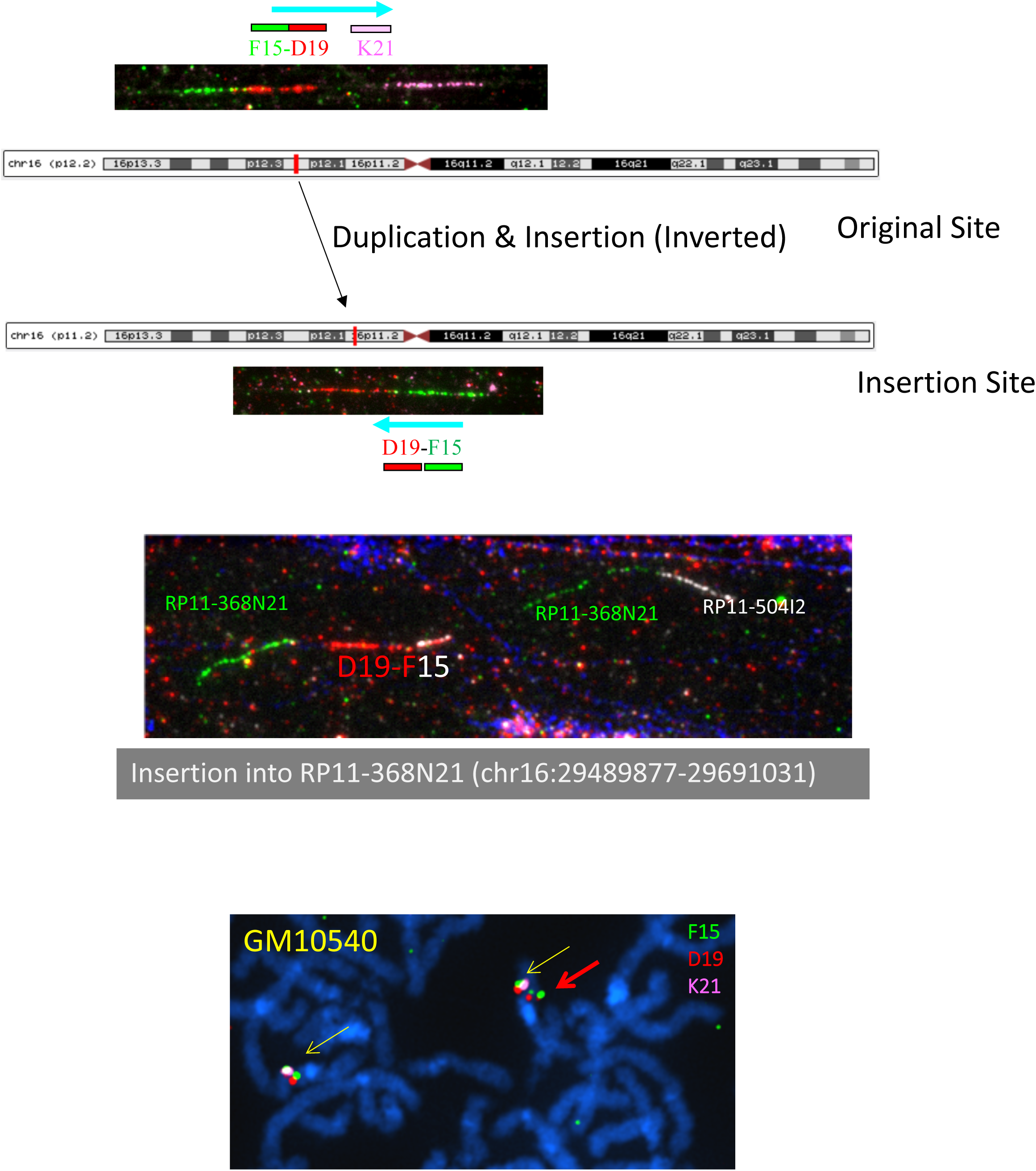
fibre-FISH of chr16 Oceanian-specific expansion shared with Denisovan genome at ∼82% frequency in all three Oceanian populations. **Top:** Cartoon illustration of location of original (16p12.2) and inserted site 7Mb away (16p11.2). **Centre**: Insertion (red) into region 7Mb away in clone RP11-368N21 (green). **Bottom:** Fiber-FISH of heterozygous duplication observed in metaphase (cell-line GM10540), yellow arrows show reference and red arrow shows duplication.

**Figure S13:**
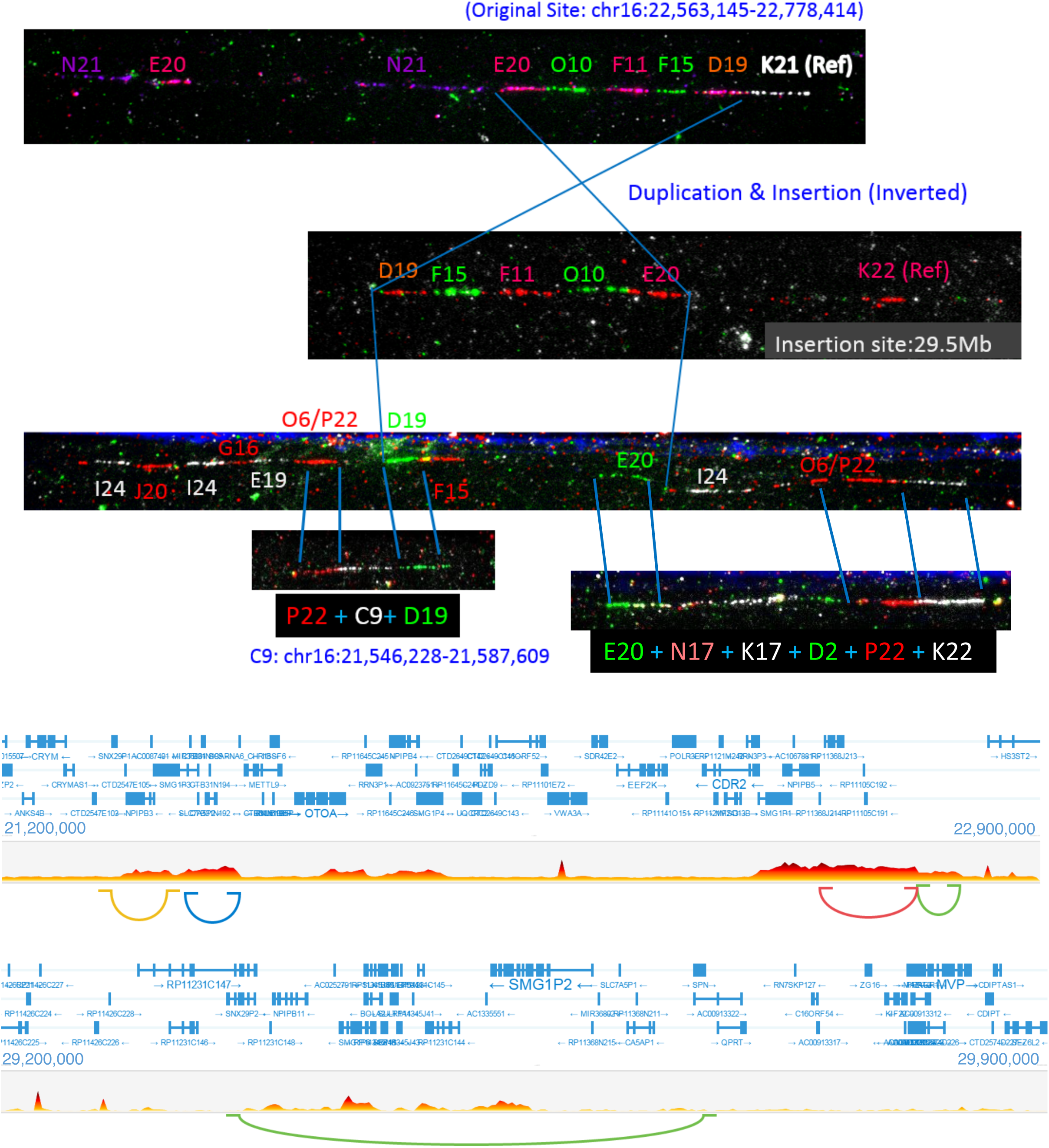
chr16p12 Papuan-specific expansion shared with Denisovan genome in more detail. **Top:** Fiber-FISH illustrating the original site (top), the (inverted) insertion sites (centre) and the region surrounding the insertion site (bottom). Region flanking the insertion site (C9) is a sequence 1Mb away from the original site, consistent with GenomeSTRiP calling a second duplication at this site in perfect LD with the initial duplication. Manta also identifies a Papuan-specific inversion at this locus. This suggests a complex event involving a duplication-inverted-insertion, an inversion and a deletion. **Bottom:** 10X-linked reads barcode overlap in region. Longranger also identifies a complex event at this locus. Top plot shows the original site barcode overlap and the regions of structural rearrangements, including the region of C9 (on the left). Bottom shows the insertion site. Note that this region is gene rich.

**Figure S14:**
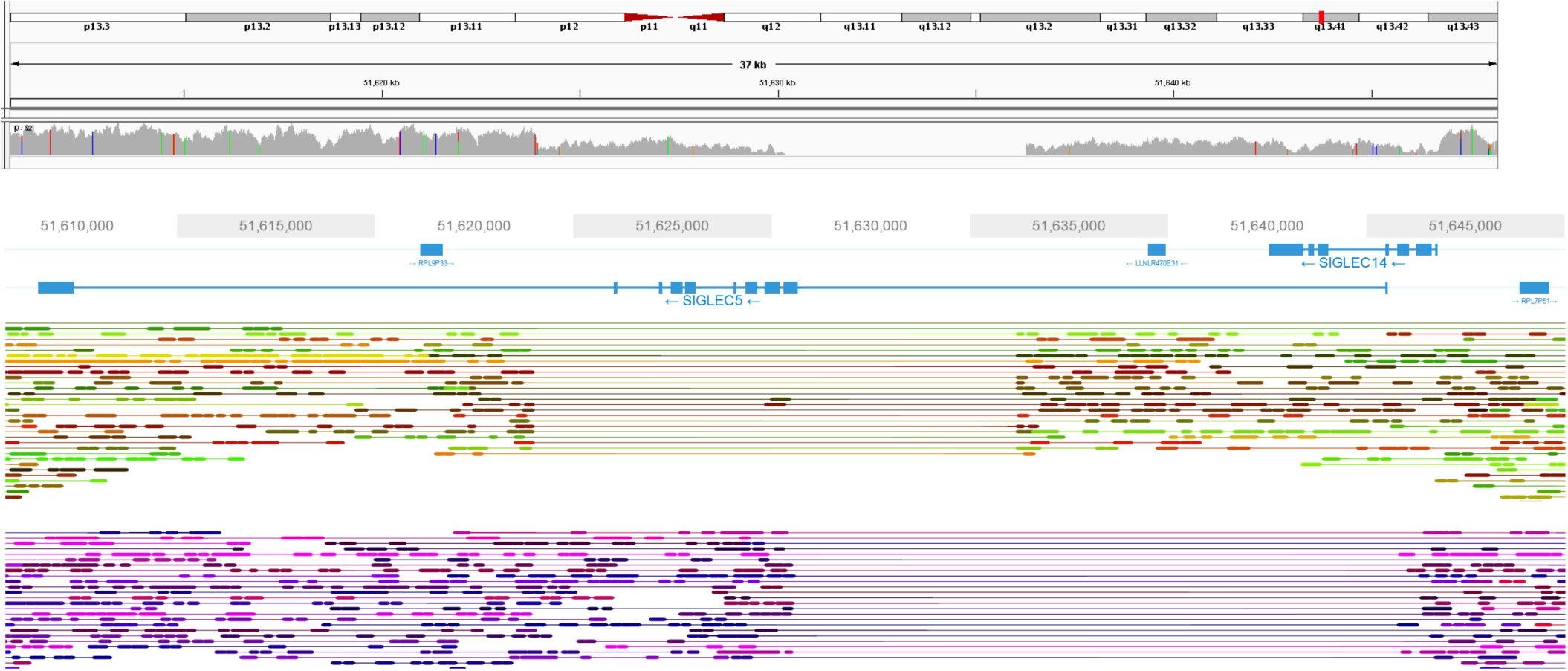
Two distinct deletions present in an Mbuti sample (HGDP00450). **Top**: IGV screenshot of depth in the region, deletions are present but appear complex. **Bottom**: Loupe screenshot of the region (as in Figure 1) showing the 2 haplotypes resolved using 10x linked-reads, each carrying a different deletion. One is the Mbuti-specific variant that deletes SIGLEC5, while the other is a common global deletion that removes SIGLEC14 creating a fused gene. Lines connecting reads illustrate that they are linked, i.e. they are from the same input DNA molecule.

**Figure S15:**
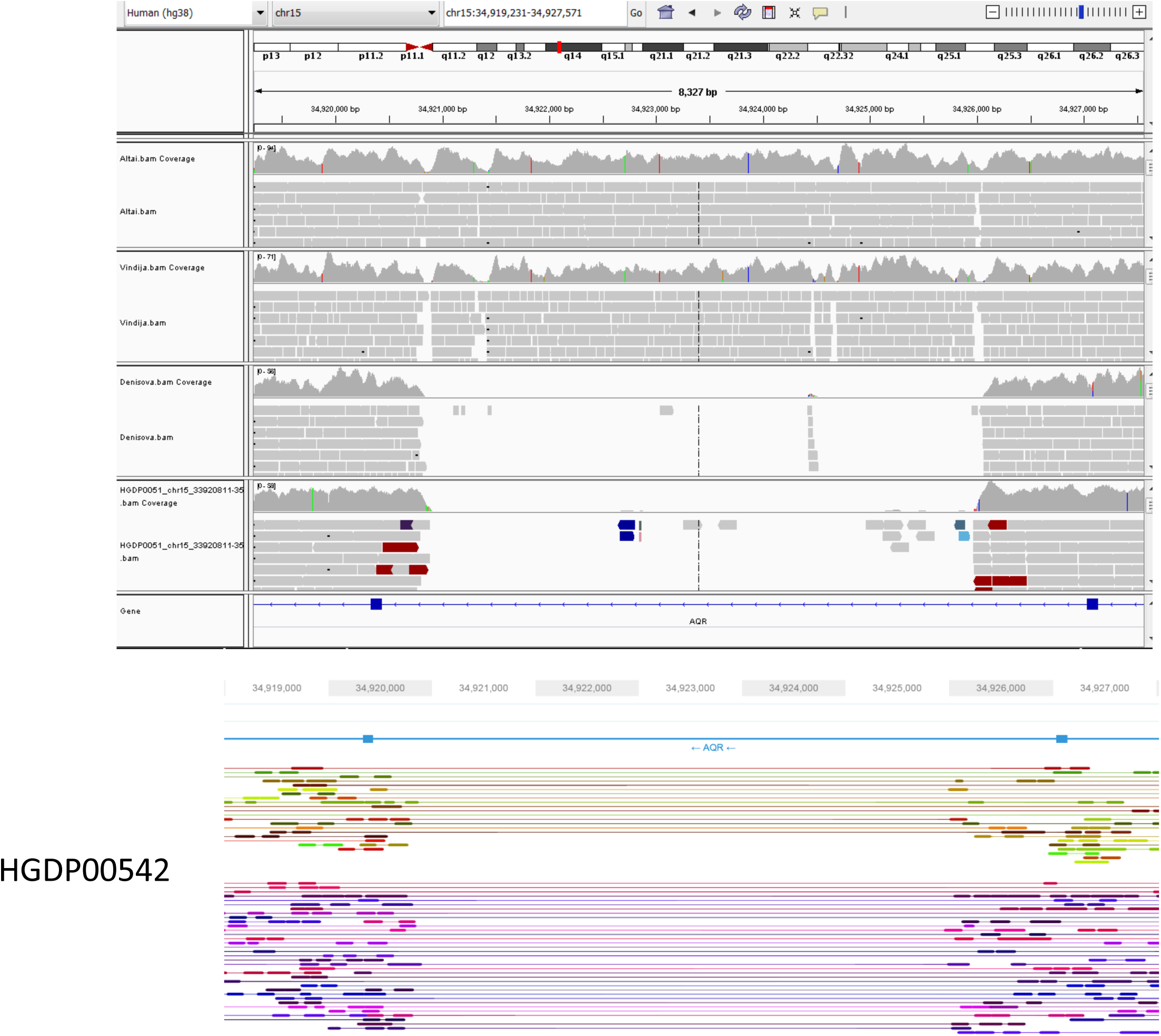
Top: IGV screenshot of a deletion in *AQR* which is present at 63% frequency in Oceanian populations. The deletion is shared with the Denisovan genome but not the Neanderthals. Also shown is HGDP00551 in the bottom IGV track (homozygous for the deletion). **Bottom**: Loupe screenshot of the region in HGDP00542 showing the 2 haplotypes resolved using 10x linked-reads, each carrying the deletion.

**Figure S16:**
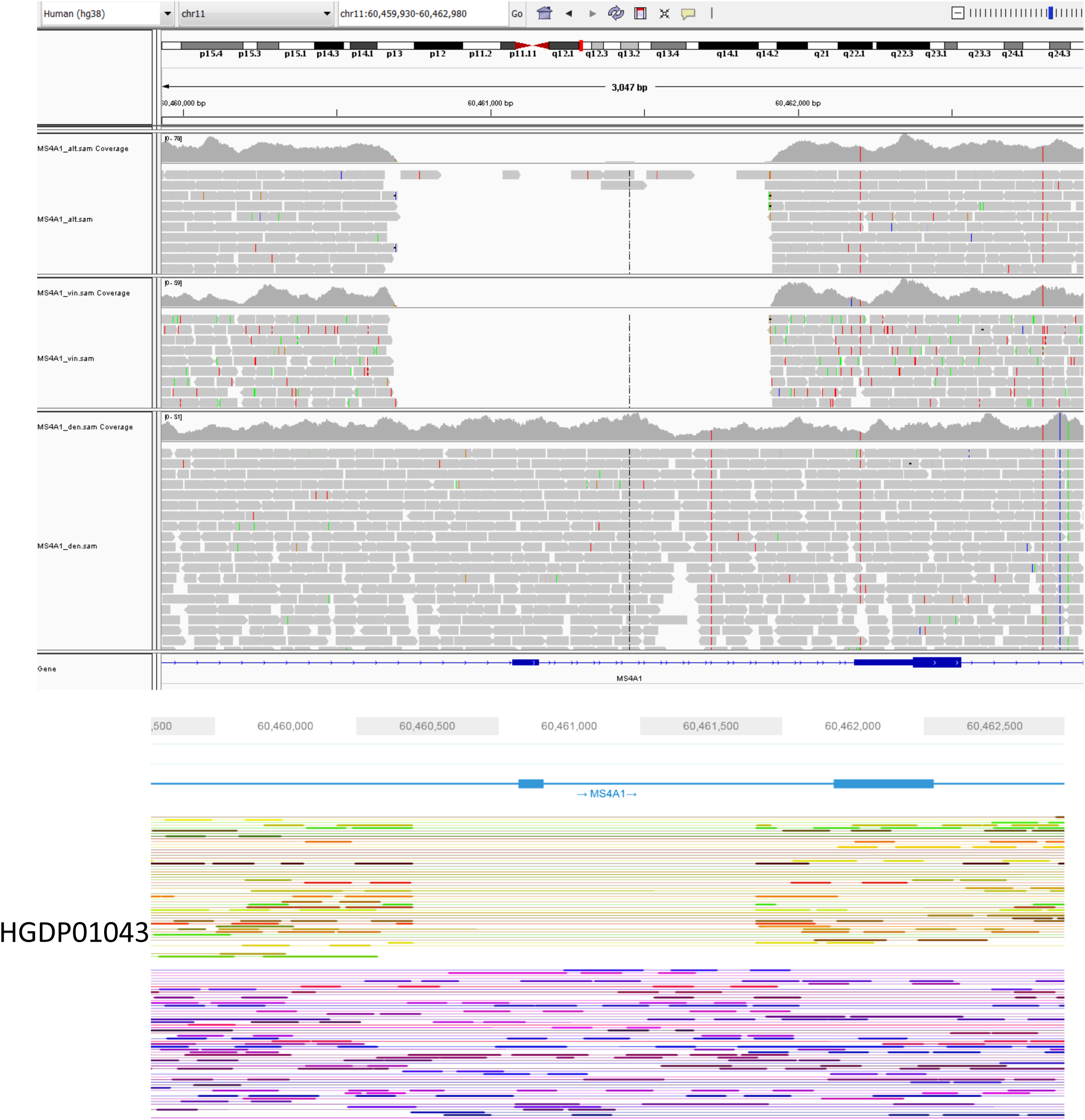
Top: IGV screenshot of a deletion in an exon of *MS4A1,* which encodes the B-cell differentiation antigen CD20. The deletion is shared by both Neanderthals (Altai top, Vindija middle track) and American populations (reaches ∼26% in Surui and Pima). The deletion is not present in the Denisovan genome (bottom track). **Bottom**: Loupe screenshot of the region in HGDP01043 showing the two haplotypes resolved using 10x linked-reads, with one carrying the deletion.

**Figure S17:**
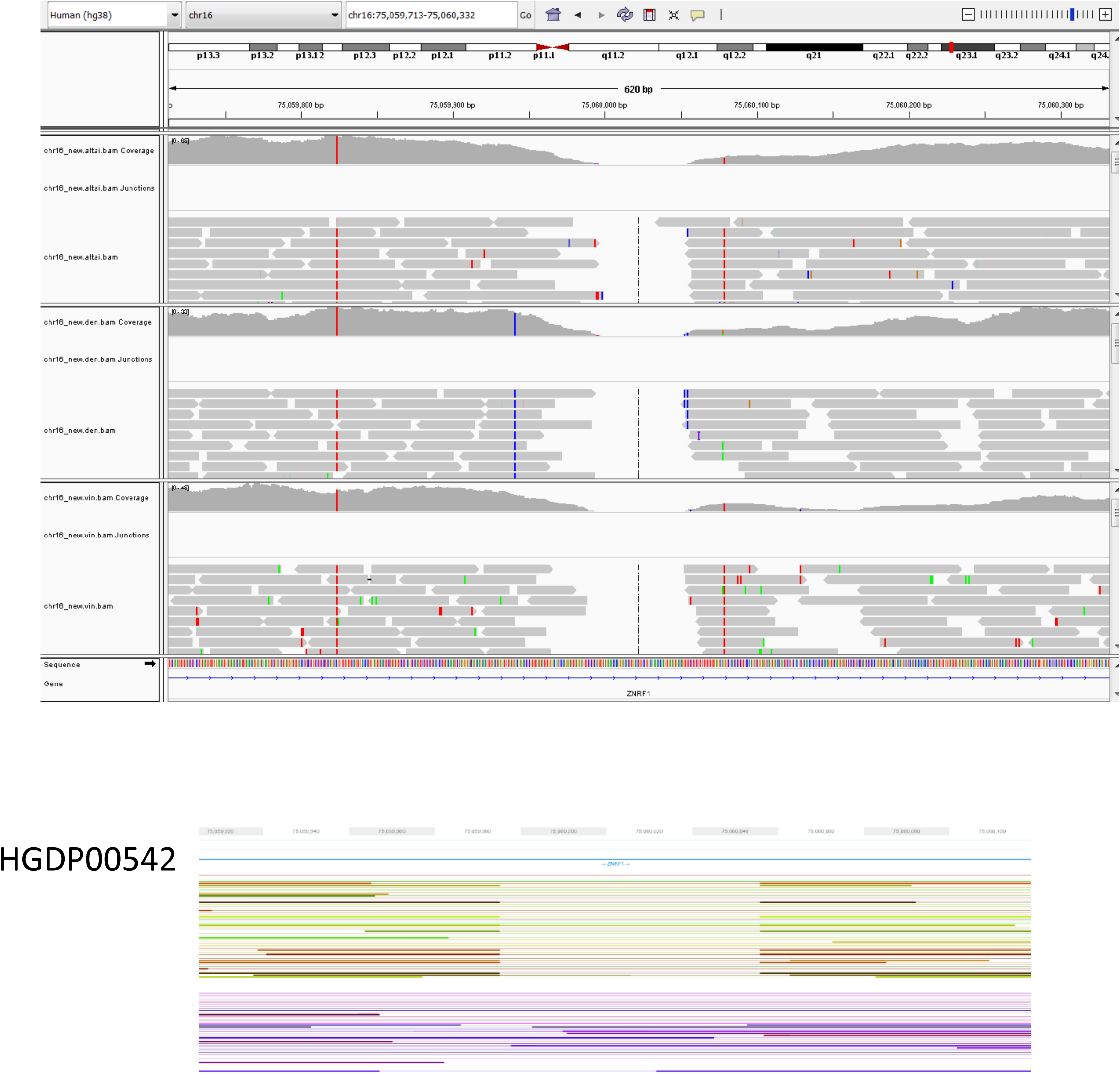
Top: IGV screenshot of a small deletion (63 bp) in *ZNRF1* which is present at 34% frequency in Oceanian populations. Top track Altai Neanderthal, middle track Altai Denisova, bottom track Vindija Neanderthal. The deletion is present in all 3 archaic genomes. **Bottom**: Loupe screenshot of the region in HGDP00542 showing the two haplotypes resolved using 10x linked-reads, with one carrying the deletion.

**Figure S18:**
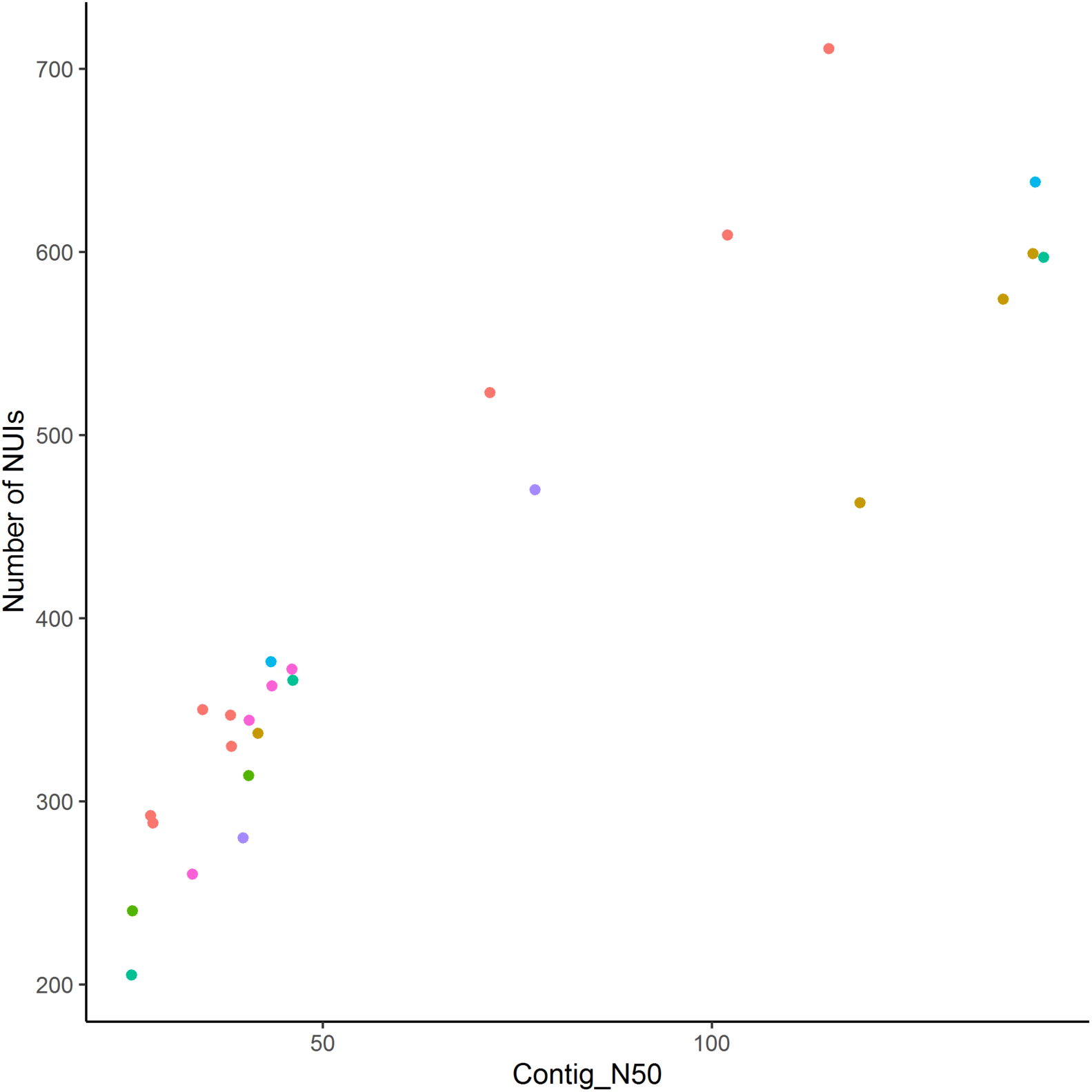
Correlation between Contig N50 and Number of identified NUIs (r = 0.91). Colours refer to the regional group of the samples.

**Table S1:**
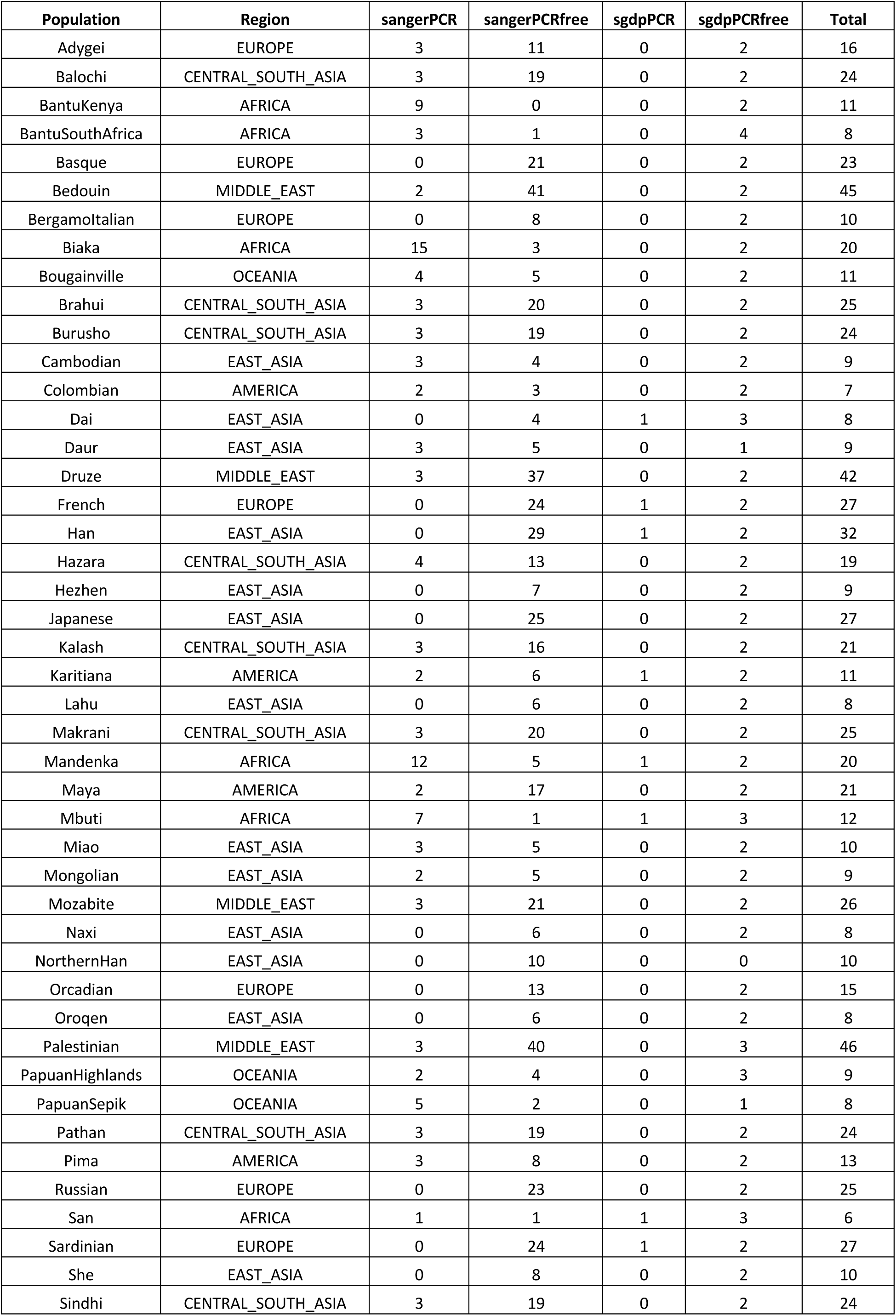
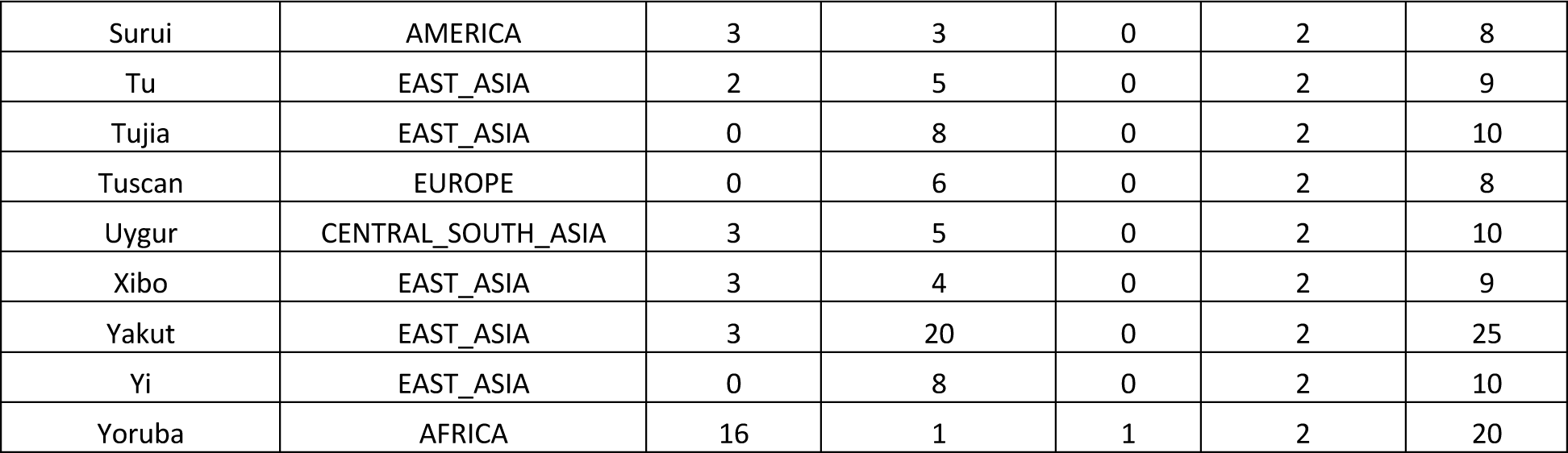
Number of samples per population analysed in this study (passing QC) stratified by library preparation and sequencing location. Total 911 samples from 54 populations.

**Table S2:** Assembly statistics and number of identified NUIs per sample.

| ID | Population | Assembly size (Mb) | ScaffN50 (Kb) | ContigN50 (Kb) | Raw_coverage | Number_NUI |
| --- | --- | --- | --- | --- | --- | --- |
| HGDP00224 | Pathan | 2110 | 32.2 | 25.6 | 52.0x | 240 |
| HGDP00228 | Pathan | 2520 | 46.6 | 40.5 | 53.4x | 314 |
| HGDP00450 | Mbuti | 2710 | 234.9 | 71.5 | 51.6x | 523 |
| HGDP00460 | Biaka | 2760 | 761.5 | 102 | 52.1x | 609 |
| HGDP00472 | Biaka | 2460 | 45.6 | 38.2 | 45.9x | 347 |
| HGDP00542 | PapuanSepik | 2510 | 68.4 | 46.1 | 52.5x | 372 |
| HGDP00547 | PapuanSepik | 2510 | 48.4 | 40.6 | 42.4x | 344 |
| HGDP00549 | PapuanHighlands | 2350 | 46 | 33.3 | 52.4x | 260 |
| HGDP00551 | PapuanHighlands | 2460 | 64.3 | 43.5 | 40.2x | 363 |
| HGDP00562 | Druze | 2500 | 47.2 | 39.8 | 50.8x | 280 |
| HGDP00580 | Druze | 2730 | 303.7 | 77.3 | 46.8x | 470 |
| HGDP00670 | Sardinian | 2510 | 68.7 | 43.4 | 48.1x | 376 |
| HGDP00774 | Han | 2490 | 70.3 | 46.2 | 49.9x | 366 |
| HGDP00819 | Han | 2790 | 8330 | 142.6 | 52.4x | 597 |
| HGDP00930 | Yoruba | 2150 | 34.3 | 27.9 | 46.1x | 292 |
| HGDP00931 | Yoruba | 2410 | 46 | 38.3 | 48.4x | 330 |
| HGDP00946 | Yakut | 2040 | 30.9 | 25.5 | 47.7x | 205 |
| HGDP01013 | Karitiana | 2790 | 11160 | 137.4 | 51.1x | 574 |
| HGDP01019 | Karitiana | 2560 | 48.3 | 41.7 | 54.5x | 337 |
| HGDP01029 | San | 2100 | 42.1 | 28.2 | 45.7x | 288 |
| HGDP01032 | San | 2760 | 8480 | 115 | 48.9x | 711 |
| HGDP01043 | Pima | 2790 | 7370 | 141.2 | 50.9x | 599 |
| HGDP01056 | Pima | 2790 | 15440 | 119 | 54.0x | 463 |
| HGDP01067 | Sardinian | 2780 | 16310 | 141.5 | 52.4x | 638 |
| HGDP01081 | Mbuti | 2260 | 51 | 34.6 | 47.6x | 350 |

**Table S3:** List of clones that mapped to the location of the chr16p12 Oceanian-specific duplication shared with the Denisovan genome. Probes were designed using the hg18 reference. Lifted over position in GRCh38 are provided.

| Probe Name in Figures | Probe ID | Chr16 Position (hg18) | Lifted over Position (GRCh38) | Note |
| --- | --- | --- | --- | --- |
| C9 | WI2-1834C9 | 21465050-21506431 | chr16:21546228-21587609 | Original Site |
| D2 | WI2-431D2 | 22359681-22400825 | chr16:22440859-22482003 | 5' Reference (Also Segmental Duplication at insertion site) |
| K17 | WI2-555K17 | 22409260-22452683 | chr16:22490438-22533861 | 5' Reference (Also Segmental Duplication at insertion site) |
| N17 | WI2-1747N17 | 22464671-22508847 | chr16:22545849-22590025 | 5' Reference (Also Segmental Duplication at insertion site) |
| E20 | WI2-3914E20 | 22481967-22524149 | chr16:22563145-22605327 | Original Site |
| O10 | WI2-916O10 | 22526829-22568957 | chr16:22608007-22650135 | Original Site |
| F11 | WI2-810F11 | 22584142-22624353 | chr16:22665320-22705531 | Original Site |
| F15 | WI2-1829F15 | 22625840-22662216 | chr16:22707018-22743394 | Original Site |
| D19 | WI2-2529D19 | 22654207-22697236 | chr16:22735385-22778414 | Original Site |
| K21 | WI2-694K21 | 22727030-22769938 | chr16:22808208-22851116 | 3' Reference |
| N21 | RP11-368N21 | 29408699-29609853 | chr16:29489877-29691031 | Insertion site |
| J20 | WI2-2731J20 | 29369385-29408106 | chr16:29450563-29489284 | Insertion site |
| I24 | WI2-3063I24 | 29414050-29452497 | chr16:29495228-29533675 | Insertion site |
| G16 | WI2-1802G16 | 29446690-29485090 | chr16:29527868-29566268 | Insertion site |
| E19 | WI2-3518E19 | 29469634-29511102 | chr16:29550812-29592280 | Insertion site |
| O6 | WI2-0456O6 | 29510348-29548504 | chr16:29591526-29629682 | Insertion site |
| P22 | WI2-1399P22 | 29528833-29568651 | chr16:29610011-29649829 | Insertion site |
| K22 | WI2-2372K22 | 29559162-29602429 | chr16:29640340-29683607 | 3' Reference |
| I2 | RP11-504I2 | 29609848-29784210 | chr16:29691026-29865388 | 3' Reference |

